# Eusociality and the evolution of aging in superorganisms

**DOI:** 10.1101/2021.05.06.442925

**Authors:** Boris H. Kramer, G. Sander van Doorn, Babak M. S. Arani, Ido Pen

**Author notes:** The authors wish to be identified to the reviewers.

## Abstract

Eusocial insects – ants, bees, wasps and termites – are being recognized as model organisms to unravel the evolutionary paradox of aging for two reasons: (1) queens (and kings, in termites) of social insects outlive similar sized solitary insects by up to several orders of magnitude; (2) all eusocial taxa show a divergence of long queen and shorter worker lifespans, despite their shared genomes and even under risk-free laboratory environments. Traditionally, these observations have been explained by invoking classical evolutionary aging theory: well-protected inside their nests, queens are much less exposed to external hazards than foraging workers, and this provides natural selection the opportunity to favor queens that perform well at advanced ages. Although quite plausible, these verbal arguments have not been backed up by mathematical analysis. Here, for the first time, we provide quantitative models for the evolution of caste-specific aging patterns. We show that caste-specific mortality risks are in general neither sufficient nor necessary to explain the evolutionary divergence in lifespan between queens and workers and the extraordinary queen lifespans. Reproductive monopolization and the delayed production of sexual offspring in highly social colonies lead natural selection to inherently favor queens that live much longer than workers, even when exposed to the same external hazards. Factors that reduce a colony’s reproductive skew, such as polygyny and worker reproduction, tend to reduce the evolutionary divergence in lifespan between queens and workers. Caste-specific extrinsic hazards also affect lifespan divergence but to a much smaller extent than reproductive monopolization.

## Introduction

The evolution of eusociality represents the latest major evolutionary transition (or transition in individuality) and has shaped co-evolved behavioral, demographic and ecological traits via an interplay between society, the individual and the environment (Johnson and Carey 2014; Michod 2000; Quiñones and Pen 2017; Smith and Szathmary 1997). Paradoxically, life in eusocial colonies seems to violate one of the cornerstones of life history theory – the trade-off between individual survival and fecundity (Kramer et al. 2015; Negroni et al. 2016) – since queens of eusocial species typically are both highly fecund and long-lived, with some ant and termite queens having lifespans of multiple decades. In contrast, workers in social insect colonies are often sterile and have much shorter lives than their queens (Fig. 1) (Hölldobler and Wilson 1990; Keller and Genoud 1997; Kramer and Schaible 2013a). Strikingly, this lifespan divergence of queens and workers does not reflect genetic differences between the castes, but emerges from differential gene expression patterns in identical genomes (Evans and Wheeler 1999; Hoffman and Goodisman 2007). Such uniquely large intraspecific variation in the rate of aging makes eusocial insects ideal organisms for studying the proximate and ultimate causes of aging.

**Figure 1:**
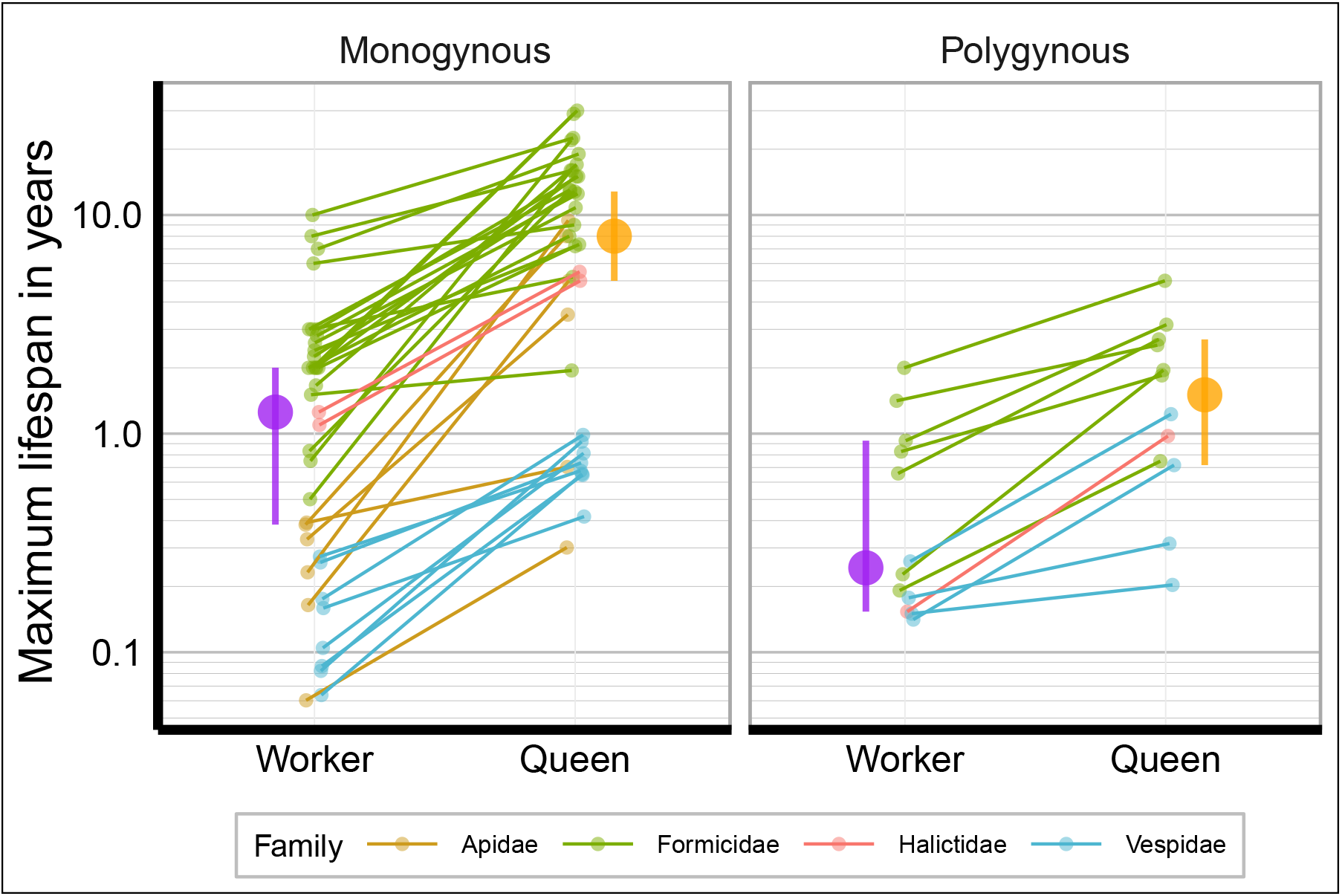
Divergence of queen and worker lifespans in eusocial Hymenoptera. Differently colored lines indicates families and connect queens and workers of the same species, purple (orange) dots show the median worker (queens) lifespans and the corresponding 0.95 confidence intervals. The left panel shows monogynous species (*n* = 37, median queen to worker lifespan ratio = 5.0 (CI: 4.0 – 7.8), median queen lifespan = 8.0 (CI: 5.0 – 12.8), median worker lifespan = 1.3 (CI: 0.4 – 2.0)) and the right panel shows poygynous species (*n* = 12, median lifespan ratio = 3.7 (CI: 1.8 – 5.1), median queen lifespan = 1.5 (CI: 0.7 – 2.7), median worker lifespan = 0.24 (CI: 0.15 – 0.93)). Phylogenetic ANOVAs show that queens have shorter lifespans in polygynous species than in monogynous species (*F* = 9.578, *p* = 0.0167) while there is no difference for workers (*F* = 3.165, *p* = 0.1614) or the lifespan ratios (*F* = 3.854, *p* = 0.1204). Data taken from Kramer and Schaible 2013a, species used can be found in Table S1.

The classical evolutionary theories of aging (Kirkwood 1977; Medawar 1952; Williams 1957) are based on the notion of a declining “force of selection” with age (Hamilton 1966), i.e. natural selection is less effective at weeding out deleterious mutations that are harmful only late in life than at removing mutations that are harmful in earlier life. The logic is that late-acting mutations are more likely to be passed on to offspring before their effects are materialized, or they are never expressed at all because their carriers die beforehand. Thus, if selection is weak enough, late-acting harmful mutations can become fixed by genetic drift and accumulate over evolutionary time, leading to decreased intrinsic rates of survival and fecundity in older individuals (Medawar 1952). If such mutations also have pleiotropic positive effects at younger ages, then natural selection can actively assist their spread to fixation despite their negative effects at older ages (Williams 1957). Based on a declining force of selection, an important prediction of the classical evolutionary theories of aging is that an increase in “extrinsic” mortality, such as predation and disease, promotes the evolution of more rapid aging, provided the increased mortality disproportionately lowers the frequency of older individuals and density dependence does mainly affect survival or fertility late in life (Abrams 1993; Kramer et al. 2016b). Thus, species exposed to higher rates of extrinsic mortality should age more rapidly than less exposed species.

Building on this predicted role of extrinsic mortality, it has been argued for over 20 years that the evolved differences in lifespan between queens and workers are readily explained by the classical evolutionary theories of aging (Keller and Genoud 1997; Parker 2010; Rueppell et al. 2015). Since social insect colonies act as “fortresses” in which queens are protected at the center, while workers are exposed to extrinsic hazards when foraging or defending the nest, the force of selection should decline with age faster in workers than in queens, hence the observed divergence of queen and worker lifespan. While on lifespans in social insects is scarce (Keller and Genoud 1997; Kramer and Schaible 2013a), information on age-specific mortality is very rare (Kramer et al. 2016a). This may be the reason why formal demographic analysis on the evolution of aging in social insects has barely been carried out.

Although prima facie not implausible, theoretically it is far from clear whether caste-specific extrinsic mortality is necessary or even sufficient to explain the divergence in lifespans between queens and workers, or if other factors such as the social structure may be more important (Giraldo and Traniello 2014). Indeed, plausible arguments can be made to suggest that the classical models will fail when applied to social insects. The classical aging models assume purely age-structured populations and therefore do not conform to the peculiar biology of social insects with their caste-structure and reproductive division of labor (Amdam and Page 2005; Kramer et al. 2016b), indeed it has been shown that Hamilton’s predictions can differ when implementing the complex interplay between age and size, stage or caste (Colchero and Schaible 2014; Steiner et al. 2014). Furthermore, the classical models assume that all offspring have the potential to, sooner or later, become reproductive themselves, while in eusocial insects most offspring (the workers) never reproduce, and therefore have zero reproductive value. This may have two important effects on the evolution of caste-specific aging. First, according to the classical models, the force of selection is zero for age classes with zero reproductive value, such as post-reproductive age classes, which implies that workers should face extremely rapid aging from birth onwards. However, workers do gain indirect fitness benefits from helping their mother raise additional siblings, and models that allow for menopausal resource transfer between mother and offspring (Lee 2003) suggest that this might delay the onset of aging in workers, although it is not clear by how much. Secondly, queens usually start producing reproductive offspring, future queens and males, after an initial phase during which they only produce workers. Thus, compared to solitary breeders with similar age-schedules of reproduction, queens typically delay the production of offspring with non-zero reproductive value (Hölldobler and Wilson 1990; Kramer et al. 2016b). According to the classical models, the age-specific force of selection is maximal and invariant with respect to age before the onset of first reproduction (Hamilton 1966; Kramer et al. 2016b). This suggests that for queens the force of selection is also maximal before the onset of producing reproductive offspring, and this may go a long way in explaining why queens live longer than otherwise comparable solitary insects. Note that this argument holds independent of reduced extrinsic mortality afforded by a defensible nest, suggesting that caste-specific extrinsic mortality is not necessary for the evolution of long lifespans in queens and the divergence in lifespans between social insect castes. In addition, it is important to consider the social structure of eusocial organisms; while most species have only one reproductive queen in their colonies (monogyny), some species display polygyny, a derived trait where colonies have multiple queens (Boomsma et al. 2014; Hughes et al. 2008). Polygyny is associated with shorter queen lifespans when compared to monogynous species (Fig. 1) (Keller 1998; Keller and Genoud 1997). Again, it has been suggested that differences in extrinsic mortality between monogynous and polygynous queens drive the lifespan differences, but the ultimate causes for shorter queen lifespans in secondary polygynous species remain unresolved (Boomsma et al. 2014; Keller and Genoud 1997). An alternative hypothesis for shorter lifespans of polygynous queens can be based on the view that colonies are superorganisms. From this perspective, the number of queens in a colony has strong implications for the evolution of aging: the death of the single queen in a monogynous society with sterile workers will lead to the death of the colony including the loss of its future reproductive success. If selection acts on whole colonies, queen survival and fertility are the factors that maximize a colonies fitness, while selection on queen lifespan and fertility should be relaxed in polygynous species because queen death does not lead to the death of the whole colony or superorganism.

To investigate these verbal arguments more rigorously, we present mathematical and simulation models for the evolution of caste-specific aging patterns in eusocial insects. Our models explore the force of selection, simulate mutation accumulation and study the effects of social structure on the evolution of queen and worker lifespans in eusocial insects.

## Models and results

First we present an analytical model and derive expressions for the strength of selection against age-specific increases in mortality for queens and workers. Then we show results of an individual-based simulation model where we allow caste- and age-specific mutations to accumulate by drift and selection and thus investigate the long-term evolution of caste-specific patterns of mortality and longevity.

### Analytical model

We work with a continuous-time model of a large population of social insect colonies, each of which is founded by a single queen. The model is hierarchical in the sense that there is a component of between-colony dynamics and between-colony age structure of queens, determined by age-specific mortality and fecundity of queens, and a component of within-colony dynamics and age-structure of workers, determined by age-specific mortality of workers and the production of new workers by the colony. See Al-Khafaji et al. (2009) and Bateman et al. (2018) for alternative approaches. For simplicity we assume that the mortality of queens only depends on their age, and not on the size of their worker force, while the queens’ reproductive output (new workers and reproductives) only depends on the (dynamic) number of workers in the colony. We relax this assumption later in the individual-based simulation model.

We use a similar approach as Hamilton (1966) to quantify how changes in age-specific mortality of queens affect the growth rate of a population of colonies in which the age distribution of queens has reached a demographic equilibrium. However, at the within-colony level, it is not safe to assume that the age distribution of workers reaches a demographic equilibrium; a colony’s queen may die long before the age distribution of workers has stabilized. Therefore, we developed a non-equilibrium approach to model how the age structure or workers changes with colony age, using nonlinear integral equations.

We will use the variable *a*_q_ ≥ 0 to denote the age of queens, but for convenience we will also refer to *a*_q_ as the age of the colonies founded by their queens, making a mental note that queens will actually be a bit older than the colonies they founded. Likewise, the variable *a*_w_ will be used to denote the age of workers within colonies, with the obvious constraint that workers can never be older than their queens: 0 ≤ *a_w_* ≤ *a_q_*. Although in the real world the number of workers in a colony can be quite small – especially in young colonies – and therefore a stochastic model would be more realistic, for simplicity we use continuous deterministic models for the within-colony dynamics of the number of workers (but see our stochastic simulation model in the next section). To that end, let *n* (*a*_w_, *a*_q_) be the number of workers of age *a*_w_ in a colony of age *a*_q_. The total number of workers in a colony of age *a*_q_ is then

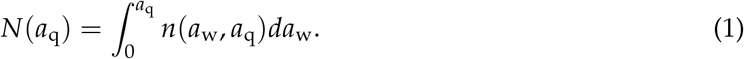

The number of offspring produced by the queen at age *a*_q_, provided she is alive, of course, is assumed to have the form

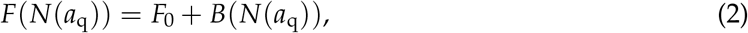

where *F*_0_ > 0 is a constant and the “benefit” *B*(*N*) is an increasing (*B’* (*N*) > 0) and bounded function with *B*(0) = 0. Thus, at colony age *a*_q_ = 0, when a queen has no workers yet (*N*(0) = 0), she produces *F_0_* offspring on her own. Let 0 ≤ *W*(*a*_q_) ≤ 1 describe the proportion of workers among the queen’s offspring produced at age *a*_q_, while *R*(*a*_q_) = 1 – *W*(*a*_q_) represents the complementary proportion of reproductive offspring, i.e. the potential future queens and males. Typically, the first offspring produced in a colony are all workers, i.e. *W*(0) = 1 (and therefore *R*(0) = 0). In older colonies *W*(*a*_q_ > 0) might behave like a (quasi) periodic function if the colony produces reproductives according to a seasonal or annual pattern (as in the example in Fig. 2).

**Figure 2:**
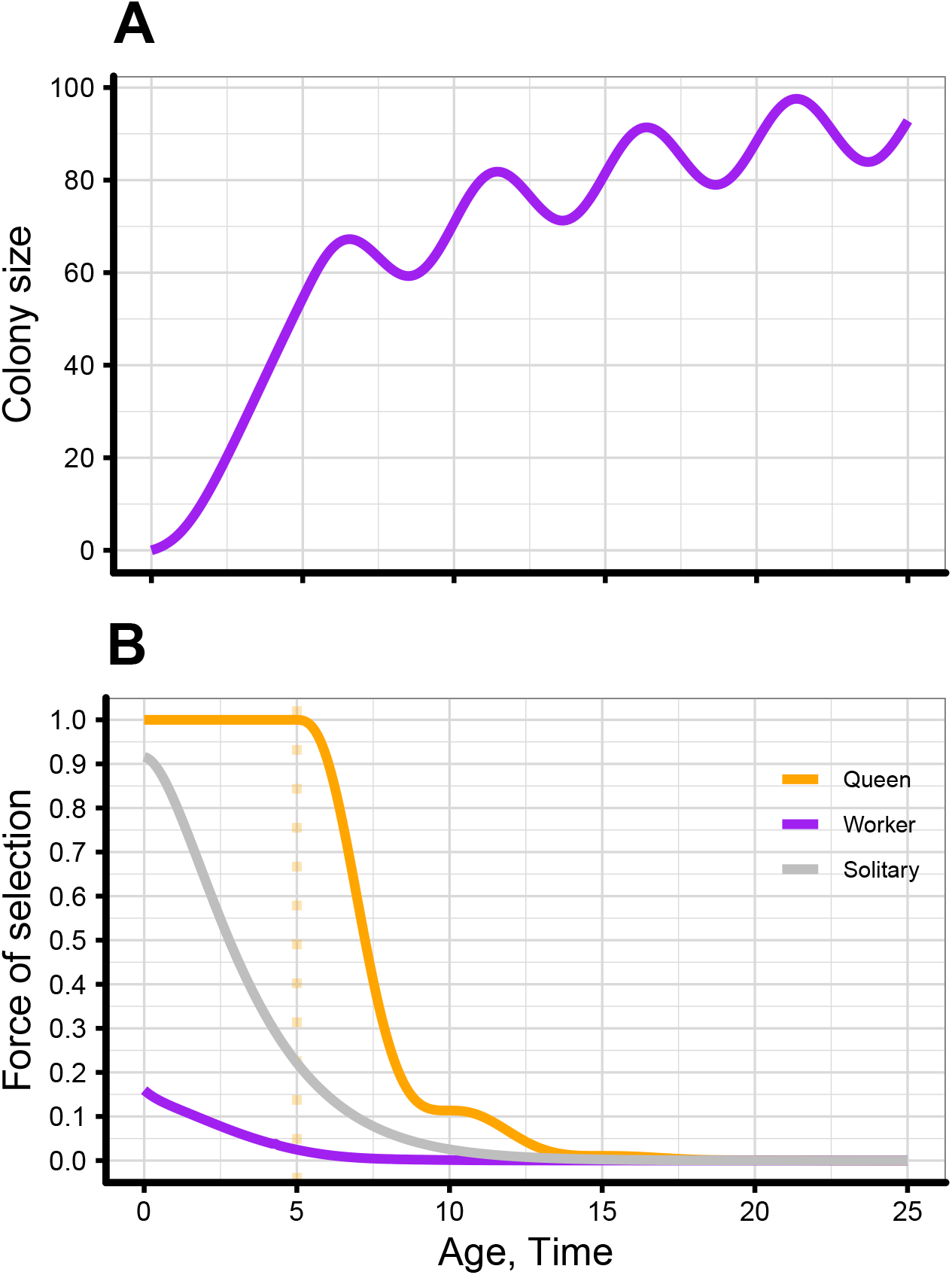
**A**, dynamics of the number of workers in a colony (eqn. 5). The fluctuations are due to the quasi-periodic production of workers: the proportion of workers among offspring is *W*(*a*_q_) = 1 for colony age *a*_q_ < α = 5 and 1 – *sin*(*a*_q_π/5)^2^ for *a*_q_ > *α*. **B**, Age-specific strength of selection for queens, workers and solitary breeders (absolute values, normalized to 1 at age 0 for queens). Queens start producing workers at colony age *a*_q_ = 0 and also reproductives at *a*_q_ = *α* = 5. Solitary breeders start producing offspring at the same age that queens start producing workers. Age-specific instantaneous mortality rates are the same for all types, *μ* = 0.1. Colony productivity as a function of the number of workers *N* is given by 2 + 20*N*/(10 + *N*). Note that selection in queens does not start to drop until the age at first production of reproductive offspring, while for workers and solitary breeders the decline starts from the get-go.

Therefore, the number of newborn workers (with age *a*_w_ = 0) at colony age *a*_q_ is given by

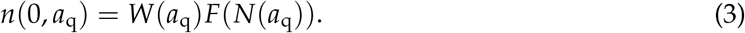

Let the age-specific instantaneous mortality rate of workers be given by *μ*_w_(*a*_w_); then their probability to survive until age *a*_w_ is given by

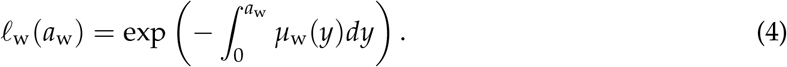

This is just the solution to the differential equation *dℓ*_w_(*a*_w_)/*da*_w_ = –*μ*_w_(*a*_w_)*ℓ*_w_(*a*_w_) with initial condition *ℓ*_w_ (0) = 1. The total number of workers *N* then obeys the inhomogeneous and typically nonlinear Volterra integral equation (VIE):

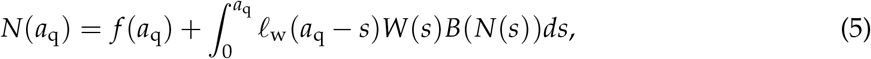

where the forcing term 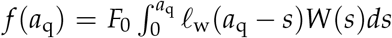. This VIE is impossible to solve analytically for *N*(*a*_q_), at least for typical nonlinear functions *B*(*N*), but under mild assumptions on *B*(*N*) the existence of unique and bounded solutions is guaranteed and it’s possible to derive some useful results for small perturbations of the solutions (see Appendix A).

Queens have their own age-specific instantaneous mortality rate *μ*_q_ (*a*_q_) and survival function

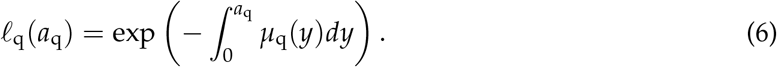

The long-term growth rate *r* of the population of colonies in demographic equilibrium, when the population-wide age distribution of queens has stabilized, is then the real solution of the Euler-Lotka equation

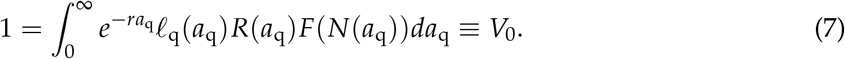

Note that the right-hand side of the Euler-Lotka is equal to a queen’s reproductive value at birth, *V*_0_ (Fisher 1930).

The potentially complex dynamics of the number of workers, governed by the VIE (5), and possibly quasiperiodic production of reproductives, governed by *R*(*a*_q_), may make *R*(*a*_q_)*F*(*N*(*a*_q_)) a complex function of colony age, which raises the question whether the population of colonies will settle down on a stable age distribution, and hence the validity of using the Euler-Lotka equation. However, the validity is guaranteed if the function is continuous and piecewise smooth (Kot 2001). Even though in reality production of reproductives can be discontinuous to some extent, we simply assume that *R*(*a*_q_) can be sufficiently well approximated by a continuous and piecewise smooth function. In the examples of Fig. 2 this is certainly the case.

We are going to apply small perturbations to the mortality functions *μ*_w_(*a*) and *μ*_q_(*a*) and investigate how this changes the growth rate *r*. Specifically, we perturb the mortality functions pointwise at age *x* by adding a delta function centered around *x* (Caswell 2019), such that after perturbation a mortality function *μ*(*a*) becomes

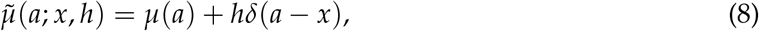

where *δ*(*a* – *x*) is the Dirac delta function or unit impulse function; this is a generalized function – a “spike” at *x* – whose most useful property here is that 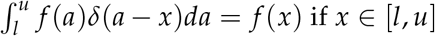 and zero otherwise. The sensitivity of a function *F*(*μ*(*a*)) with respect to a perturbation of mortality at age *x* is then defined by

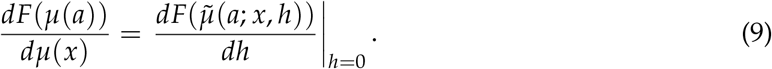

For example, the sensitivity of 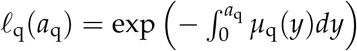 with respect to *μ*_q_ (*x*) is given by

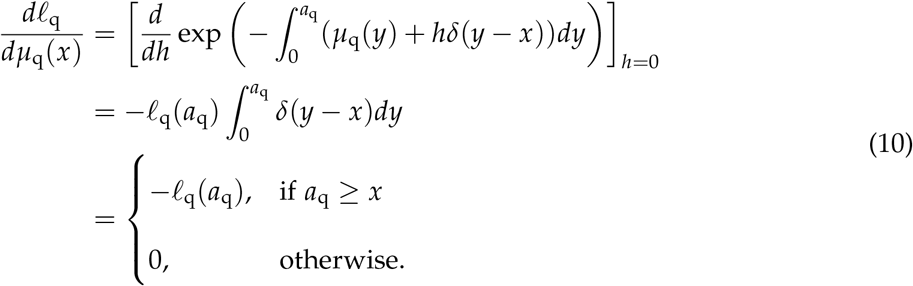

#### Selection differential for queens

We follow the implicit function approach of Hamilton (1966), applied to the right-hand side *V*_0_ of the Euler-Lotka equation (7). The selection differential for queens with respect to mortality at age *a*_q_ = *x* then equals

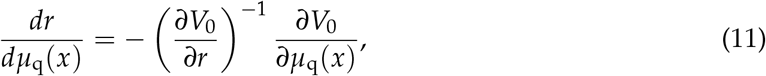

where

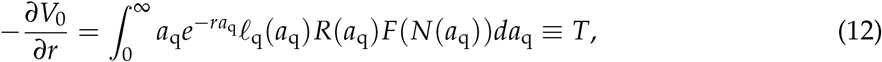

and

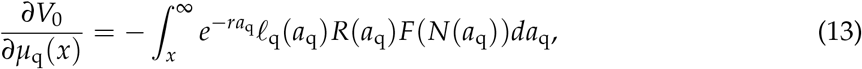

where we have used result (10) to compute the second partial derivative.

Therefore,

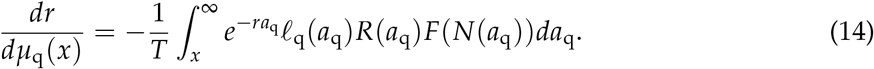

This is basically the same result as obtained by Hamilton (1966) with slightly different methods. The denominator is the mean age at which queens produce reproductives, or, in other words, the generation time *T*. It is also interesting to note that maximization of *r* is equivalent is equivalent to maximization of the right-hand side of the Euler-Lotka equation, which is the expected reproductive value at birth *V*_0_ of queens, which is just their expected number of offspring produced over a lifetime, with future offspring discounted for population growth. Expression (14) as a whole also has a nice interpretation in terms of reproductive values, analogous to a result reported for a discrete time model by Kramer et al. (2016b). We can rewrite (14) more condensely as

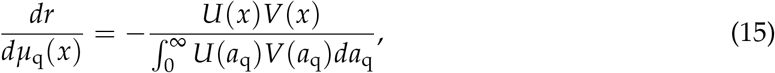

where 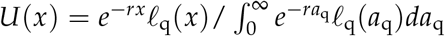 is the stable age distribution of queens, and 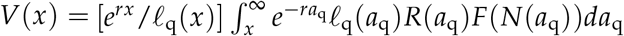 is the individual reproductive value of a queen with age *x*. The product 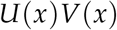 is the “class reproductive value” (Taylor 1990) of all queens with age *x*, while 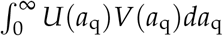 can be regarded as the population average individual reproductive value of queens, or the total class reproductive value of all queens. Thus, the selection differential for queens with respect to mortality rate at age *x* can be interpreted as minus the proportion of total reproductive value represented by all queens of age *x*.

Clearly, *dr*/*dμ*_q_(*x*) is a non-decreasing function of *x*, since the integrand in the numerator of (14) is positive and independent of *x*. Moreover, if no reproductives are produced by queens before the age of *α*, then *dr*/*dμq*(*x*) is minimal and constant until *x* = *α*, and increases for *x* > *α*, which is similar to the classical result that selection is strongest and constant until the age of first reproduction, and declines afterwards. However, in the case of social insect colonies, it is not the age of first reproduction, but rather the age at which the first reproductives are produced (Fig. 2). Reproduction in the form of workers typically precedes the first emergence of reproductive offspring.

#### Selection differential for workers

Worker mortality only enters the Euler-Lotka equation through the fecundity function *F*(*N*(*a*_q_)), which depends on the number of workers *N*(*a*_q_) present at colony age *a*_q_. Thus, when perturbing the workers’ mortality function *μ*_w_(*a*_w_) at age *x*, the resulting sensitivity of *r* can be written as

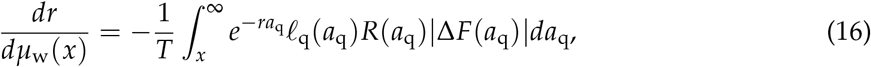

where *T* is given by (12) and Δ*F*(*a*_q_) = *F’*(*N*(*a*_q_))*dN*(*a*_q_)/*dμ*_w_(*x*). Here *dN*(*a*_q_)/*dμ*_w_(*x*) is the change in the number of workers at colonies with age *a*_q_, following the perturbation in the workers’ mortality at age *x* < *a*_q_. The lower limit *x* on the integral is due to the fact that workers of age *x* can only be present in colonies at least as old. This selection differential can also be written in terms of the stable age distribution and reproductive values:

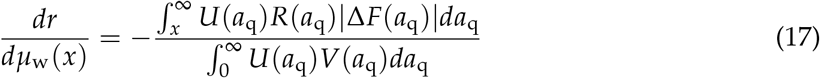

Here the numerator represents the loss in offspring (with average reproductive value *V*_0_ = 1) in colonies of all ages *a*_q_ ≥ *x*, weighted by the age class density 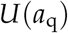, due to the loss of workers with age *x*.

In general, it is not possible to find a closed-form expression for *dN*(*a*_q_)/*dμ*_w_(*x*), but in Appendix A we show that it is always negative and increases with *x*:

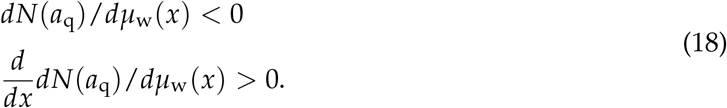

As a consequence, the strength of selection against increased worker mortality declines with worker age; however, while for queens the strength of selection does not start to decline until after the first production of reproductives, the strength of selection for workers starts to decline from birth onward (i.e. when *x* = 0) (Fig. 2). Technically, the reason is that in the expression (14) for queens the integral is independent of *x* when *x* < *α*, the age of first production of reproductives, while in the expression (16) for workers, the integral decreases with *x* even if *x* < *α* because the factor |Δ*F*(*a*_q_)| in the integrand is a decreasing function of *x*, as follows directly from (18). Intuitively, the loss in fitness to the queen is always the same (and maximal) at whatever age she dies before the colony produces its first reproductive offspring, but for workers it’s a different story: the negative effect on future colony size of increased age-specific worker mortality decreases with worker age even before the colony produces its first reproductive offspring.

Moreover, noting the similarity between the expressions (14) and (16), it is quite obvious that selection in workers is always weaker than selection in queens. This is because |Δ*F*(*a*_q_)| < *F*(*a*_q_) for any positive increasing continuous function *F*. In fact, |Δ*F*(*a*_q_)| will often be considerably smaller than *F*(*a*_q_) for reasonable choices of *F*(*a*_q_). Intuitively, the loss of a queen at age *x* implies the full loss of the reproductive value of a queen of age *x*, while the loss of a cohort of workers of age *x* will only cause a partial loss of reproductive value of the queen.

Strictly speaking, the caste-specific selection differentials derived above apply to the differential spread of mutant alleles that affect age-specific mortality in asexual populations. However, we are mainly interested in the relative strengths of selection between castes, and we argue that adjusting the model to encompass sexual reproduction and/or different genetic ploidy levels will not make a difference in that regard. As we have seen, the selection differentials for queens (eqns. 14 and 15) and for workers (eqns. 16 and 17) represent a reduction in the reproductive value at birth of queens, a reduction in future offspring production due to increased mortality of queens or workers. In order to account for sexual reproduction and differing ploidy levels, we can view the selection differentials as inclusive fitness effects, where we must weight the lost offspring by their relatedness to the “actor” class, i.e. the class in which the mortality-enhancing alleles are expressed, and by their sex-specific class reproductive values (Taylor 1988). Thus, we can expand the fecundity term *F* as follows:

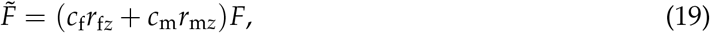

where *c*_f_ and *c*_m_ refer to class reproductive values of female and male offspring, respectively, while *r_fz_* and *r_fz_* to the relatedness of female and male offspring to the actor class *z*, either queens (*z* = *q*) or workers (*z* = *w*). The class reproductive values can be further expanded in terms of offspring sex ratios (proportion sons *s*) and sex-specific individual reproductive values (*υ*_f_ and *υ*_m_): *c*_f_ = (1 — *s*)*υ*_f_ and *c*_m_ = *sυ*_m_. Sex-specific individual reproductive values, in turn, depend on ploidy and the sex ratio: when both sexes are diploid 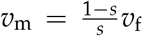, while in haplodiploids 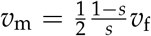. The factor (1 — *s*) / *s* represents the expected number of mates per male, while the extra factor 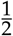 in haplodiploids reflects the fact that in haplodiploid species males only pass on genes via daughters while females transmit them via offspring of both sexes. Putting all this together:

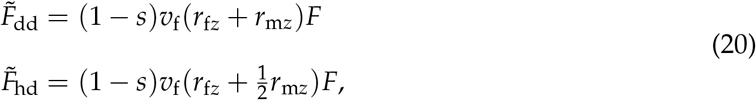

where the subscripts dd and hd refer to diplodiploidy and haplodiploidy, respectively. For diplodiploids we further have 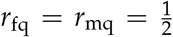 and 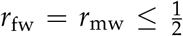 where for workers the upper bound holds for singly-mated queens. For haplodiploids, 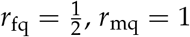 and 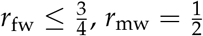. Plugging the relatedness coefficients for species with singly mated queens into (20) yields

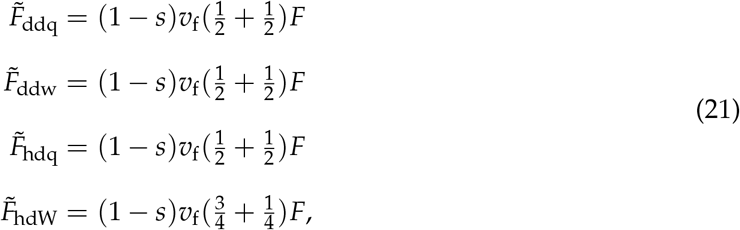

which are all identical to (1 — *s*)*F* in demographic equilibrium when *υ*_f_ = *V*_0_ = 1. Thus, for species with singly mated queens, sexual reproduction, varying sex ratios and varying ploidy do not affect the relative strengths of selection between castes. For species with polyandrous queens, selection is relatively weaker in workers because workers are more distantly related than queens to the female offspring they help raise. Also note that the absolute strength of selection is weaker under sexual reproduction by a factor (1 — *s*), which is the proportion of daughters among reproductive offspring. This also males sense, because we assume that mortality-increasing alleles are only expressed in females.

Figure 2 shows some numerical examples, where queens, workers and solitary breeders have exactly the same age-specific mortality rates (i.e. no differences in “extrinsic” mortality), yet the strength of selection against higher mortality is much stronger in queens, which predicts that queens should age much more slowly than workers and solitary breeders. In the example, both queens and solitary breeders start producing offspring at the same age, but queens produce only workers for a while, which explains why in queens the strength of selection starts to decline later than in solitary breeders.

### Individual-based stochastic simulation model

We support the predictions of the analytical model with more realistic stochastic individual-based simulations by incorporating the complex interplay between social structure (monogyny, polygyny, colony inheritance), caste-specific extrinsic mortality, and worker reproductive potential on the evolution of caste-specific aging patterns (details in Appendix B). In contrast to this level of socio-demographic complexity, we stick to the simplest of theoretical frameworks regarding aging mechanisms – the mutation accumulation theory (Medawar 1952). Colonies (*N*_col_ = 1000, Table 1) in our simulations start with a single queen and no workers. As long as the number of workers is small, queens need to spend part of the time foraging in order to obtain enough resources for raising workers; once the queen produces enough workers (≈ 5) the queen only produces eggs, using the resources that workers, which all act as foragers (if they are sterile or the colony is queenright) bring back to the colony. Extrinsic mortality only acts on individuals that leave the colony. New queens and males are only produced if colonies die and need to be replaced due to colony death in the simulation. Individuals in the model are haplodiploid and their intrinsic age-specific survival probabilities are genetically encoded by gene loci with independent age- and caste-specific effects that are subject to mutations. To implement the mutation accumulation theory (Medawar 1952), mutations are on average biased towards lower survival (mutation rate, *m* = 0.001; mutation bias *η* = 0.2, see Table 1). Actual survival probabilities in the simulations are determined by multiplying intrinsic (caste- and age-specific) and extrinsic survival probabilities (only for individuals that leave the colony). At the start of each simulation, all age classes (we generally used *ω* = 20, but see SI) have the same high intrinsic survival probability, resulting in an exponentially declining cumulative survival curve with age. Figure 3 shows a typical example of the initial and evolved caste-specific survival curves after up to 20,000 time steps. In all simulations, both in queens and workers, selection removes deleterious mutations and increases intrinsic survival up to a certain threshold age, after which survival rapidly drops to zero, resembling a “wall of death” (Wachter et al. 2013). As a result baseline mortality in all simulations is low for both castes as indicated by the slowly declining survival curves at early age-classes (Fig. 3). Aging rates (increase in age-specific mortality) are also similar for both as indicated in Figure 3, but the onset of aging - which drive the differences in lifespan and life expectancy in the simulations - occurs at earlier ages in workers as predicted by the analytical model. We therefore refer to lifespans for the rest of the manuscript.

**Figure 3:**
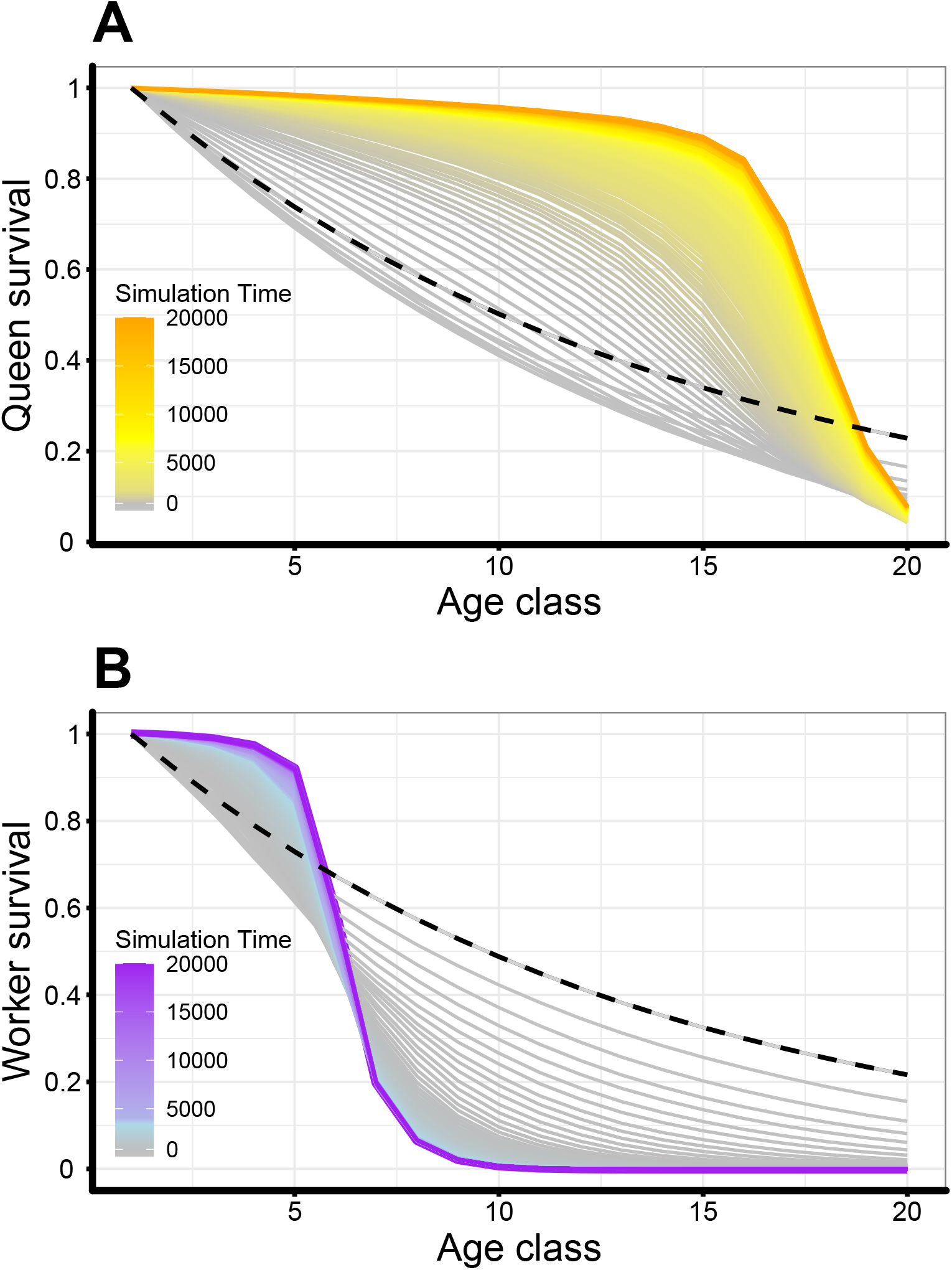
Initial (dashed) and evolved (solid) survival curves for queens (A) and workers (B). Depicted are means of 50 simulation replicates for the monogyny scenario with sterile workers. Parameter settings: *h_e_* = 0, *r* = 1.0, *f_q_* = 5.0, *f_w_* = 0,

**Table 1:** Parameters and parameter value ranges for the individual-based simulation models.

| Parameter | Explanation |
| --- | --- |
| $N_{\text{col}}$ | Number of colonies per simulation (1000) |
| $\omega$ | Number of age classes (20) |
| $h_e$ | Extrinsic hazard. Probability of death per time step (0.0-0.3) |
| $r$ | Productivity constant. Resources per unit time per forager (1-6) |
| $f_q$ | Queen fecundity: number of eggs per unit time (1-6) |
| $f_w$ | Worker fecundity: number of eggs per unit time step (in queenless colony) (0-6) |
| $m$ | Probability of mutation per gene per reproductive event (0.001) |
| $\eta$ | Mutation bias towards higher mortality (0.2) |
| $\sigma$ | Mutational effect size (0.08) |
| $\alpha$ | Mutational partial correlation between adjacent age classes (0-0.5) |
| $\beta$ | Mutational correlation between castes (0-0.5) |

Since the number of queens or the chance of workers to inherit colonies, as seen in wasps, has been shown empirically to correlate with caste-specific lifespans (Fig. 1)(Bourke 2007; Keller and Genoud 1997; Kramer and Schaible 2013a), three different social structure scenarios were implemented: (i) Monogyny, where colonies only have one queen that cannot be replaced after her death. After queens death, the remaining workers may produce males if they are fertile until all workers have died. Once all individuals disappeared from the colony the colony is replaced by a new queen produced in one of the remaining colonies in the simulation. (ii) In the serial polygyny scenario, colonies have a single queen that can be replaced by a new queen produced in the same colony, in case of the queen’s death. (iii) Colony inheritance, where in case of the death of the queen a random worker of the colony expresses the queen phenotype, as seen in some primitively eusocial wasps. Colonies in (ii & iii) are usually queen right thus workers do not get the chance to produce offspring even if they are fertile.

In the monogyny scenario queen and worker lifespans always diverge over evolutionary time (Fig. 4A, 4B). This divergence also appears without any caste-specific extrinsic mortality, as predicted by the analytical model and is most pronounced when workers are sterile (Fig. 4A). When workers are fertile, queens’ lifespans are shorter and worker lifespans are longer as compared to the sterile case and the ratio between queen and worker lifespans increases with increasing exposure to extrinsic mortality (Fig 4A). Increased extrinsic mortality has a slightly negative effect on the evolved queen-worker lifespan ratio when workers were sterile, as queens may get exposed to extrinsic mortality when colonies are small due to worker loss, while the effect is positive and stronger when workers were fertile (Fig. 4B). In both cases queen lifespan decreases at the highest extrinsic mortality setting, because the declining workforce exposes the queen to extrinsic mortality.

**Figure 4:**
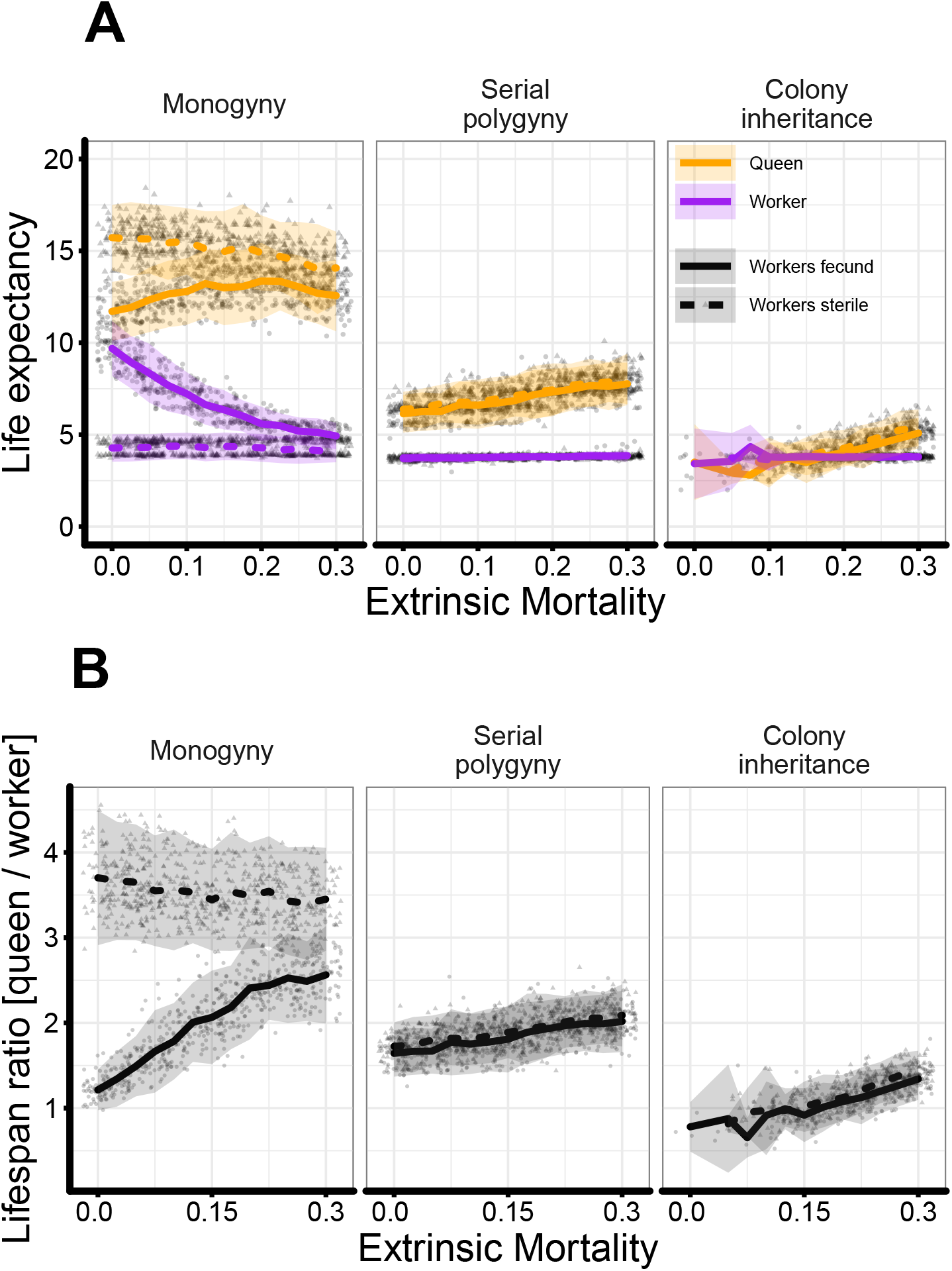
Effects of extrinsic mortality and social structure on the evolution of queen and worker lifespans. Grey dots indicate mean evolved values from individual simulation runs; lines represent means of 50 simulations with fecund (solid) and sterile (dashed) workers; shaded areas represent two standard deviations from the mean. Panel A shows evolved queen and worker life expectancies at birth, panel B shows the corresponding queen to worker lifespan ratios. Parameter values: *r* = 2, *f_q_* = 3.0, *f_w_* = 0 or 3.0, *α* = *β* = 0.

In the serial polygyny scenario colonies are potentially immortal as lost queens can be replaced by a queen produced in the same colony. All simulations implementing the serial polygyny scenario show the evolution of diverging queen and worker lifespans but the evolved lifespan ratios are considerably smaller than in the monogynous scenario with sterile workers (Fig. 4B). Without extrinsic mortality queen lifespans are on average 50% longer than worker lifespans and with increasing extrinsic mortality queen lifespan increases across all parameter settings (see supplemental material) while worker lifespan show only a negligible response (Fig. 4A, 4B, S1, S2). Queen lifespans are generally shorter than in the monogyny scenario when no extrinsic mortality was added to the model but show an increase in lifespan alongside increasing extrinsic mortality. Since colonies are potentially immortal and nearly always queenright, workers do not get the chance to reproduce. Consequently worker fertility shows no effect on the evolved lifespans and colony sizes were larger at high levels of extrinsic mortality than in the monogynous scenario (Fig. S1).

Across all parameter settings, the colony inheritance scenario leads to the evolution of the shortest queen and worker lifespans. Differences between queen and worker lifespan are solely driven by increasing extrinsic mortality (Fig. 4, S1, S2). Queen lifespans double with increasing risk while worker lifespan shows negligible changes (Fig. 4B, S1, S2). Since colonies are always queenright worker fertility does not show an effect.

## Discussion

It has been argued for the last 20 years, that due to caste-specific differences in exposure to extrinsic sources of mortality, the classical evolutionary theories of aging readily explain the divergence of queen and worker lifespans, without formal application to social insects. Here we have shown that such differences in extrinsic mortality between queens and workers are in general neither necessary nor sufficient to account for the caste differences in lifespan but that reproductive division of labor is the main driver in the eusocial systems analyzed here. We used an analytical model to demonstrate that the strength of selection against increased agespecific mortality is always greater for queens than for workers, regardless of any patterns of caste-specific mortality, including, for example, higher mortality of queens. The expressions (14-15) and (16-17) we derived for the selection differentials against higher mortality at age *x* in queens and workers, respectively, clarify why this is so. The selection differential for queens is of the form — *C*_x_ / *C*_total_, where *C*_x_ is the class reproductive value of queens with age *x*, which is just the remaining lifetime reproductive success of all queens of age *x*, adjusted for population growth, while *C*_total_ is total reproductive value of all queens in the population. Thus, the selection differential for queens represents the loss of reproductive value of queens upon their death at age *x*. In contrast, the selection differential for workers has the form — *p*C*_x_* / C_total_, with 0 < *p* < 1, and it represents only a fractional reduction in the class reproductive value of queen with age *x* that results from the death of a cohort of workers of age *x*. Since a single cohort of workers represents only a fraction of the total number of workers, that together determine the reproductive capacity of a colony, the fitness consequences of higher age-specific mortality of workers are necessarily less severe than similar changes in the mortality for queens.

The analytical model also points to a further cause that enhances the strength of selection in queens relative to that in workers; namely the fact that the strength of selection for queens is constant and maximal until the colony produces its first reproductive offspring, i.e. the future queens and males. This is, of course, similar to Hamilton’s result for solitary breeders (Hamilton 1966), that selection is constant and strongest until the age of maturity, after which it declines. However, queens, during the ergonomic phase of somatic colony growth, typically do not produce any offspring with reproductive value – that is, offspring with the capability of transmitting copies of their mother’s genes to future generations – until the colony has reached a critical mass or favorable time of the year (Oster and Wilson 1978), which can be years after the queens produces her first (worker) offspring. Thus, one could say that for social insects it’s not the age at maturity of the queen that sets the time after which selection starts to decline in force and consequently aging rates increase, but rather the age at maturity of the colony as a whole.

In contrast to queens, where selection does not start to weaken until the onset of releasing reproductive offspring, for workers the strength of selection starts to decline from eclosion onwards (Fig. 2) pointing to an earlier onset of aging that leads to reduced lifespans. The reason is that higher age-specific mortality of workers always reduces the future number of workers, and thus the future reproductive output of the colony, and that this effect on future colony size declines with the age of workers from eclosion onwards according the quite general result in (18). Intuitively, early increased mortality has a forward-rippling effect on the numbers of workers in more age classes of workers than later increased mortality, which affects a smaller number of worker cohorts.

It has been argued before that kin selection may lead to reduced worker lifespans (Bourke 2007; Lucas and Keller 2014), but without formal models. Especially, sterile workers can only gain indirect fitness benefits by helping to raise the brood of their relatives and do so from their moment of birth, hence at an age younger than the queen or a solitary breeder, leading to a decline from birth in the force of selection as shown in Figure 2 (Bourke 2007; Lucas and Keller 2014). Although our analytical model was not formulated as a kin selection model, the expressions for the caste-specific selection differentials (14) and (16) can nevertheless be interpreted as inclusive fitness effects. The absence of coefficients of relatedness in the expressions boils down to an implicit assumption that queens and workers are equally related to the reproductive offspring produced by the colony, which is the case for monogynous colonies with singly mated queens. More generally, the selection differentials can be multiplied by caste-specific coefficients of relatedness in case queens and workers are not equally related to the reproductive offspring. This would be the case, for example, when monogynous colonies have multiply-mated queens. Workers would then be less related than queens to the reproductive offspring, which would further weaken the relative strength of selection for workers, predicting that, all else being equal, lifespan divergence between queens and workers should be even greater in monogynous polyandrous social insects, compared to monogynous monandrous species. It would be interesting to test this prediction with a comparative analysis.

The simulation model revealed that the major determinant of the evolved lifespans of queens and workers is the degree of reproductive skew which delays the onset of aging for the queen caste. The greatest lifespan divergence evolved for the monogyny scenario with sterile workers, where workers have zero chance of producing offspring, while the smallest divergence occurred when workers can be “promoted” to queens upon the death of their founding queen, as in the colony inheritance scenario (Fig. 3, 4). In general, if reproductive skew is smaller (i.e. due to more queens per nest and/or male production by workers), then elimination of a queen leads to a smaller decline in future reproductive output of the colony, and hence to weaker selection against increased queen mortality and/or stronger selection against increased worker mortality.

Even if between-caste differences in exposure to extrinsic hazard are not necessary for the evolution of caste-specific aging, as has been argued (Keller and Genoud 1997), the simulation model showed that such differences can, and often do, positively affect the evolved lifespan ratio (Fig. 4A, S1, S2), if reproduction is not restricted to a single queen and workers have the chance to reproduce. For monogynous colonies with sterile workers, increasing the exposure of workers to extrinsic mortality had almost no effect – indeed, even a slightly negative effect – on the evolved lifespan ratio, as shrinking colony sizes expose queens to extrinsic mortality. In all other scenarios we considered, the queen-worker lifespan ratio tended to increase with greater worker exposure to extrinsic mortality, but its quantitative effect tended to be relatively small compared to the effect of a colony’s social structure (Fig. 4). Extrinsic mortality may therefore have played a more significant role during early group formation and the evolution of division of labor (Hamilton 1971) as it does in systems where reproductive division of labor has already evolved, but this was not the focus of our study.

The serial polygyny scenario predicted shorter evolved queen lifespans than the monogyny scenario (Fig. 4A), consistent with results of comparative analyses of queen lifespans (Fig. 1) (Keller and Genoud 1997; Kramer and Schaible 2013a). It has been argued (Keller and Genoud 1997) that queens in polygynous colonies are more prone to extrinsic mortality and, following the logic of evolutionary aging theory, shorter queen lifespans should evolve in response. However, in the serial polygyny scenario of our simulation model queens evolved shorter lifespans not because they were more exposed to extrinsic hazards, but rather because the lifespan of a queen in a polygynous colony is of less importance for the fitness of the colony than under monogyny, where the queen’s death precipitates colony extinction. Polygyny is a derived trait, and a high risk of losing the queen should favor polygyny over monogyny (Hughes et al. 2008; Keller and Genoud 1997; Nonacs 1988). But even if polygyny is favored by extrinsic mortality, we show here that the lifespans of polygynous queens are shorter independently of the influence of extrinsic mortality. Alternatively, polygynous species often resemble r-strategists, where budding and an early onset of sexual reproduction are more efficient ways to colonize new territories. Indeed, polygyny is more common in species that frequently relocate their nests and that do not build complex and sheltered nests, relaxing selection for long queen lifespans (Hölldobler and Wilson 1990; Keller and Genoud 1997).

While missing information on age-specific mortality forced us to compare lifespans and prevented us to compare the onset of aging or aging rates with empirical data, empirical evidence shows that species with the largest lifespan divergence also show discrete morphological caste phenotypes (Kramer and Schaible 2013a). With our simulation model we also studied effects of genetic correlations between queen and worker phenotypes (the mutational covariance matrix at the genetic basis of the simulation model allows for mutations that affect the queen phenotype as well as the worker phenotype, see supplemental information). It turned out, perhaps not surprisingly, that the lifespan divergence between queens and workers increases with decreasing genetic correlations between queen and worker phenotypes (SI, Fig. S4). Thus, the independence of caste-specific phenotypes (Feldmeyer et al. 2014) appears to be a requirement for increasing differences of caste-specific lifespans. Future transcriptome studies might shed further light on these phenotypic correlations.

The most pronounced lifespan ratios in our models evolved for the monogyny scenario with sterile workers (Fig. 4B), where queens outlived workers by a factor of about four. The limited empirical data on social insects shows: the most extreme lifespan ratios of up to 30 are found among highly eusocial ant species with very large colony sizes (>10,000) and multiple phenotypically divergent worker castes which are not covered by our simulation model (Kramer and Schaible 2013a). However, median lifespan ratios in social insects are smaller than five (Fig. 1), especially in species with small colony sizes and a single worker caste, and are further reduced in primitively eusocial species where workers can become reproductives (Kramer and Schaible 2013a). One factor that is not considered in our model are the different morphological phenotypes of social insect castes that are associated with different resource costs for the production of queens and workers. These differential investments into queen and worker phenotypes may explain why our model does not capture the extreme lifespan differences seen in some ant species. In our model the production as well as maintenance costs for queens and workers are similar because our focus is on the mutation accumulation mechanism, but it has been shown that differential investment into the castes can lead to fitness benefits and the further divergence of queen and worker phenotypes and may explain even larger lifespans ratios than we find here (Jeanne 1986; Kramer and Schaible 2013b).

A “superorganism” perspective may help to think about how selection shapes caste-specific aging patterns (Amdam and Page 2005; Boomsma and Gawne 2018; Flatt et al. 2013; Kramer and Schaible 2013b). Although social insects colonies are the hallmark of animal cooperation, colonies can also be rife with conflict (Bourke 1999). The greater the reproductive skew, or concentration of reproductive value in a smaller proportion of the colony’s inhabitants, the more queens (and kings in termites) can be regarded as the “germ line” of the superorganism, while the workers embody a more-or-less disposable “soma”. If workers are sterile, reproductive division of labor leads to a mutual dependence of queen and worker phenotypes where queen lifespan is equal to colony lifespan (Boomsma and Gawne 2018). Hence the age of first reproduction of a colony is not the time it produces its first offspring, which is destined to become part of the soma, but rather the time it produces the first reproductive offspring, the progenitors of the next generation’s super-organismal germline. This is especially important when comparing monogynous and polygynous social structures: in monogynous societies, the germ line is reduced to one individual, whose survival is essential for the entire superorganism, driving the evolution of extremely long lifespans, while polygyny provides several “germ line individuals”. Consequentially, the survival of the superorganism is not based on a single queen individual, resulting in relaxed selection on queen lifespan, while creating potentially immortal colonies. In addition, the germ line-soma differentiation is linked to a trade-off between the maintenance of the “expensive” germ line versus the maintenance of the soma, which may explain why workers generally live shorter. Experiments on zebrafishes showed that the removal of the germ line leads to increased investments into the soma (Chen et al. 2020), and workers in queenless colonies may start to develop ovaries and live longer (da Silva-Matos and Garófalo 2000; Dixon et al. 2014).

Besides different levels of relatedness, complex eusocial societies can be compared to simple multicellular organisms. Recently it has been suggested that social anatomy, referring to colonies being composed of specialized individuals, and social physiology, the set of mechanisms that coordinate the activity and the development of specialized individuals, (Friedmann et al. 2020) parallel anatomy and physiology of multicellular organisms. Thus, the study of caste-specific aging and its underlying mechanisms may shed light on the evolution of aging patterns in early metazoans and on tissue-specific aging in advanced metazoans such as ourselves. This perspective might create a framework to understand the evolution of multicellular organisms by looking at the evolution of eusocial societies and vice versa. When focusing on aging, we are still lacking evolutionary explanations or predictions for relative lifespans of different tissues, and we believe that a better understanding of the different selective pressures on the evolution of different phenotypes in colonies can inform us about similar processes in unicellular organisms to achieve a more general picture on the evolution of aging in general.

## Acknowledgments

The authors would like to thank the handling editor and two anonymous reviewers for their valuable feedback that improved the manuscript. manuscript. We thank the Center for Information Technology of the University of Groningen for providing access to the Peregrine high performance computer cluster.

## Appendix A: Details of analytical model

### Uniqueness and boundedness of VIE solutions

Brauer 1972 studied inhomogeneous Volterra integral equations (VIEs) of the form

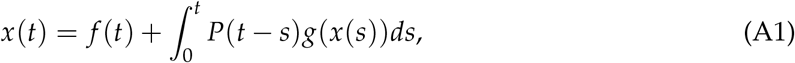

where *f* (*t*) is nonnegative, continuous and of bounded variation on *t* ≥ 0 such that lim_*t*→∞_ *f* (*t*) exist, *g*(*x*) is a nonlinear continuous nonnegative function on *x* ≥ 0 such that *g*(0) = 0 and *g’*(*x*) is continuous, *P*(*t*) is nonnegative, monotone nonincreasing and differentiable on *t* ≥ 0 and normalized such that *P*(0) = 1 and 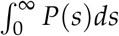 exists. Then for a given *f* (*t*) a unique nonnegative bounded solution *x*(*t*) exists, provided that

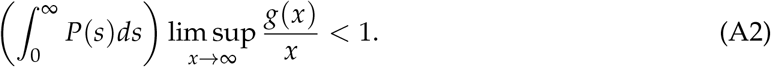

The VIE (5) is almost of the same form as (A1), with 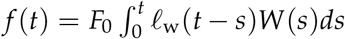 and *P*(*t*) = *ℓ*_w_(*t*), except that in our case *g*(*s, N*(*s*)) = *W*(*s*)*B*(*N*(*s*)) is a function of two variables (*s* and *N*(*s*)) rather than a single variable (*N*(*s*)) as in Brauer’s theorem. However, inspection of Brauer’s proof reveals that his theorem extends also to our case.

Let’s see if Brauer’s assumptions regarding his functions *f* (*t*), *P*(*t*) and *g*(*x*) also apply to our model. First, our *f* (*t*) is also (assumed) nonnegative and continuous. It is also of bounded variation, because since 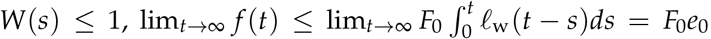, where *e*_0_ is a worker’s life expectancy at birth, which is certainly finite. Second, *ℓ*_w_(*t*) is also nonnegative, nonincreasing, differentiable and *ℓ*_w_(0) = 1. The remaining requirement for boundedness of solutions *N*(*t*) is then that 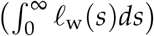 limsup_*N*→∞_ *B*(*N*)/*N* < 1. This is also true since we assume that *B*(*N*) has a finite upper bound and hence *B*(*N*)/*N* → 0 as *N* → ∞.

#### Results for the selection differential in workers

First we derive the perturbation in the number of workers when the workers’ age-dependent mortality function *μ*_w_(*a*_w_) is perturbed by a delta function centered at age *x*. Taking the derivative of (5) with respect to *μ*_w_ (*x*) gives

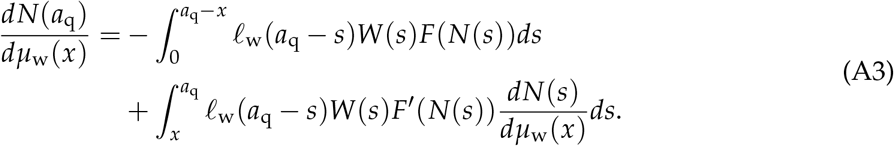

This is a linear VIE of the form

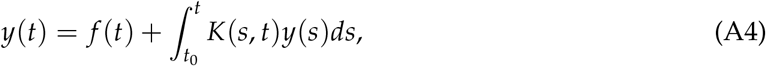

with forcing term equal to 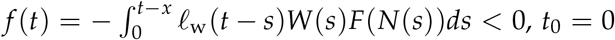 and with a kernel *K*(*s,t*) = *ℓ*_w_(*t* — *s*)*W*(*s*)*F’*(*N*(*s*)) ≥ 0. It is a classic result (e.g. Brunner 2017), going back to Volterra (1896), that the solution *y*(*t*) of a linear VIE can be expressed in terms of the so-called resolvent kernel *R*(*s, t*) as follows:

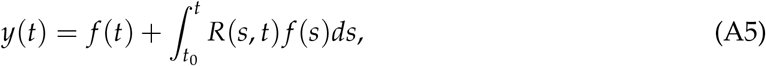

where

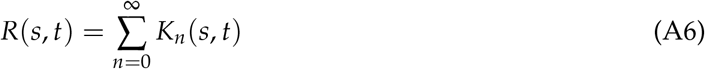

and the iterated kernels *K_n_* (*s, t*) are defined recursively as *K*_1_(*s, t*) = *K*(*s, t*), and for *n* ≥ 2

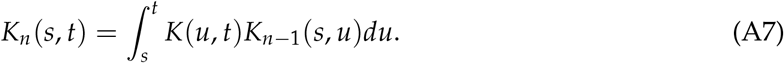

Now it is obvious that if a kernel *K*(*s, t*) is nonnegative, then so must be its associated resolvent kernel *R*(*s, t*). Therefore, by (A5), if the kernel is nonnegative, the sign of *y*(*t*) equals the sign of *f* (*t*). Hence, *dN*(*a*_q_)/*dμ*_w_(*x*) < 0.

Now we will show that *dN*(*a*_q_)/*dμ*_w_ (*x*) increases with *x*. Taking the derivative with respect to *x* of (A3),

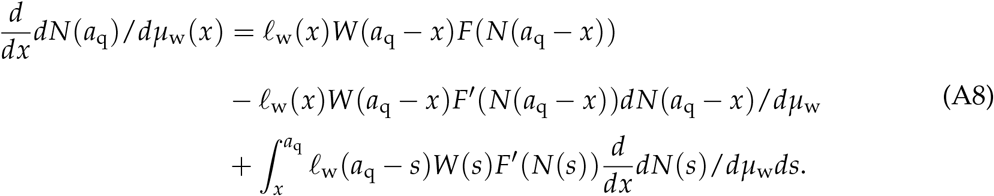

This is again a linear VIE, now with respect to (*d*/*d*_x_)*dN*(*a*_q_)/*dμ*_w_(*x*). Since the forcing term is clearly positive (noting that *F’*(*N*) > 0 and *dN*(*a*_q_)/*dμ*_w_(*x*) < 0) and the kernel nonnegative, it follows that its solution is positive, which is what we set out to show.

Comparison of the expressions for the strength of selection in queens and workers, (14) and (16), respectively, shows that selection in workers is weaker than selection in queens if |Δ*F*(*a*_q_)| = *F’*(*N*(*a*_q_))|*dN*(*a*_q_)/*dμ*_w_(*x*)| < *F*(*a*_q_). This is true because if the argument of a positive increasing function is perturbed downward, then the downward change in the function value cannot exceed the value of the function at the unperturbed argument – otherwise the function value after perturbation would be negative, which is absurd because it contradicts the assumption that the function is positive.

#### Numerical examples

In our numerical examples we use the following functions:

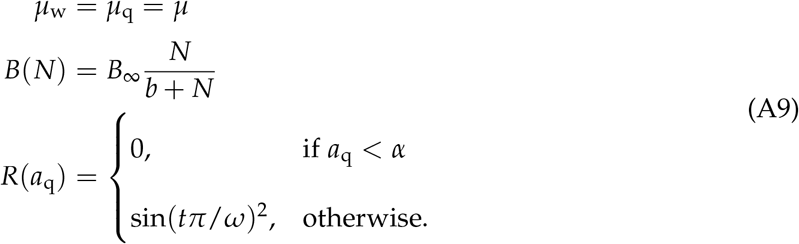

In other words, we assume that queens and workers have the same age-independent instantaneous mortality rate *μ*, that colony output due to workers increases monotonically with *N* towards an upper bound *B_∞_*, and that no reproductives are produced until colony age *a*_q_ = *α*, after which the proportion of reproductives fluctuates periodically between 0 and 1 with period *ω*.

We numerically approximated solutions of the Volterra integral equations (5) and (A3) with a simple Euler scheme. Given VIE 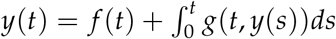, we approximated solutions with the recursion

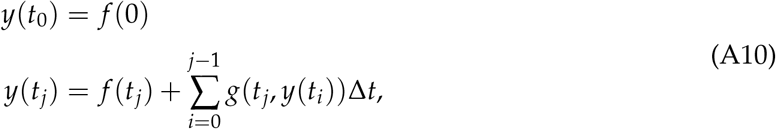

where Δ*t* = *t_j_* — *t*_*j*—1_ is a constant small time increment.

Integrals in the expressions (14) and (16) for the selection differentials of queens and workers, respectively, were numerically approximated with a similar Euler scheme.

## Appendix B: Details of simulation model

### Genetics of individual queens and workers

Each individual female starts life at age zero and has a maximum lifespan of *ω* = 20 arbitrary time units (see Table 1 for parameter values) which is determined by age-specific intrinsic and extrinsic sources of mortality. Age- and caste-specific intrinsic mortality is genetically determined by a single diploid gene locus per age class (1… *ω*) per caste, i.e. individual genomes consisted of 2*ω* = 40 diploid gene loci. Each allele is associated with a real-valued number and the two alleles at the same locus interact additively to produce a phenotypic value. Thus, the probability of intrinsic mortality between ages *x* — 1 and *x* for a member of caste *C* ∈ {*Q, W*} alive at age *x* is determined by two additively interacting alleles as follows:

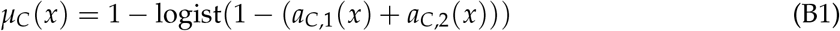

where logist(*x*) = 1/(1 + exp(—4*x*)) is a strictly increasing (logistic) function that maps the real number to the interval [0,1], and *a*_C,1_ (*x*) resp. *a*_C,2_ (*x*) are the values of maternally and paternally inherited alleles, expressed at age *x* in caste *C*. At the start of each simulation, every allele is assigned the value 0.8, implying an age-independent intrinsic survival probability of 0.96 and a corresponding negative exponential survival function (Fig. 3C, D).

### Mutation and inheritance

All genes are transmitted from parent to offspring in haploid blocks without crossing over. Prior to transmission, genomes of haploid gametes are mutated with probability *m* (typical value 0.001; see Table 1), in which case a random vector **Δ** is drawn from a 2*ω*-dimensional multivariate normal distribution and added to the allelic values of the gamete:

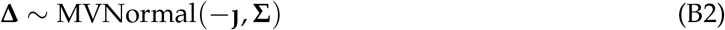

The mean mutation vector **J** < **0** is biased towards smaller allelic values (typical value —0.2; see Table 1), i.e. geared towards lower age-specific survival. For most simulations, mutations at different loci are independent, i.e. the mutational covariance matrix is of the form 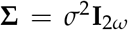, where the standard deviation *σ* is a constant “mutational effect size” (typical value 0.08; Table 1) and *I*_2ω_ an identity matrix. For a subset of simulations, we allow for mutations that are correlated across some loci; specifically, we allow for a *within-caste* positive partial correlation 0 < *α* ≤ 1 between adjacent age-classes, and for *between-caste* positive partial correlations of 0 < *β* ≤ 1 between the identical age classes of queens and workers, and a partial correlation of *β*^2^ between queens and workers of adjacent age classes. Thus, the 2*ω* × 2*ω* partial correlation matrix is of the form

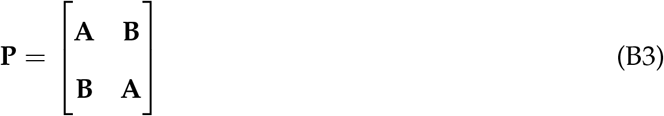

where **A** and **B** are resp. within-caste and between-caste *ω* × *ω* partial correlation matrices of tridiagonal form

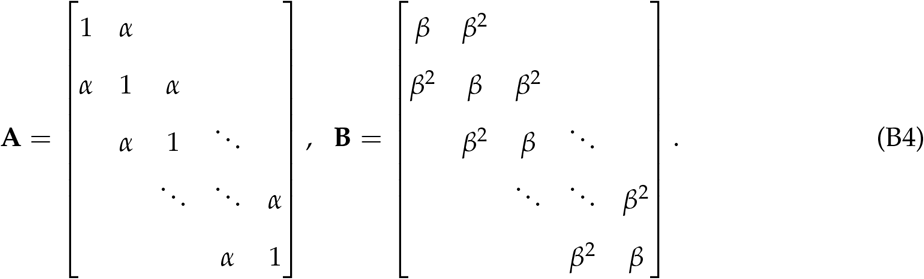

The partial correlation matrix **P** is transformed to a correlation matrix **R** according to

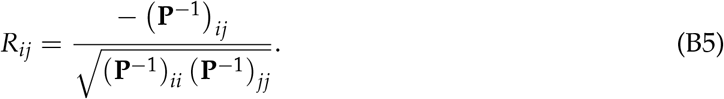

and the mutational covariance matrix is then obtained as

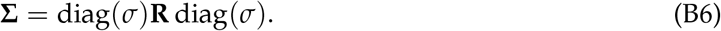

where diag(*σ*) is a diagonal matrix with constant *σ* along the diagonal.

### Foraging, fecundity, survival and colony growth

Across all simulations, each colony starts with a single queen and no workers. As long as the number of workers is small, queens need to spend part of the time foraging in order to obtain enough resources for raising workers. Specifically, let a queen spend a proportion *z* of her time foraging and 1 — *z* egg-laying. Then the number of eggs per time step is proportional to 1 — *z*,

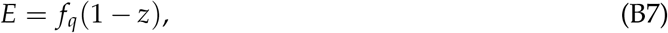

where *f*_q_ is a queen “fecundity” parameter (Table 1). The amount of resources *G* gathered per time unit is proportional to the total number of foragers, i.e. the queen a proportion *z* of the time plus the number of workers:

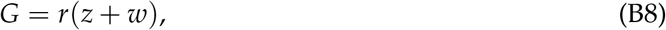

where *r* is a “productivity” constant (Table 1). We assume that the net fecundity, the number of offspring raised per time unit, is given by the product of the number of eggs and an increasing function *f* (*G/E*) (with *f’*(*x*) > 0) of the amount of resources per egg, *G/E*:

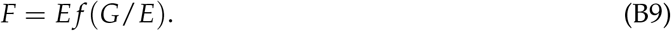

For *f* (*x*) we chose *x*/(1 + *x*) and this gives *F* = *EG*/(*E* + *G*). We assume that at every time step the queen’s allocation to foraging maximizes her net fecundity, and this implies that

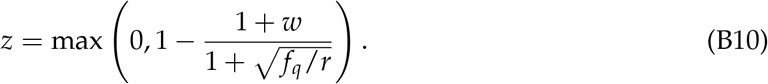

Plugging this expression into (B9), we obtain the expected brood size *F* produced by a queen per unit time, as a function of the number of workers present in the colony. In the simulations, the actual brood size *B* is a random number drawn from a Poisson distribution with parameter *F*. The resulting brood is then temporarily “stored” until the next time step, when they either joined the work force or became a reproductive male or female that establish a new colony upon extinction of another colony (see below).

At each time step every individual is potentially exposed to two hazards: it may die from “intrinsic” causes with a genetically determined probability given by (B1), and it may die from “extrinsic” causes if it spends some time foraging outside of the colony. Given a proportion of time *z* spent foraging, the probability of extrinsic mortality is given by

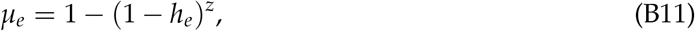

where *h_e_* is an “extrinsic hazard” constant (Table 1). Non-reproducing workers spend all of their time foraging (*z* = 1), hence their probability of dying from extrinsic causes reduces to *h_e_*, while queens with a sufficiently large worker force (B10) never forage (*z* = 0) and face zero risk of extrinsic mortality. This setup allowed us to implement increased exposure to extrinsic risk for queens when colony sizes are small.

If a queen dies, and the remaining workers in her colony are non-sterile (parameter *f_w_* > 0; Table 1), the workers can produce male (haploid) offspring on their own, facing similar time allocation trade-offs between foraging and egg-laying as queens in very small colonies. The reproductive workers’ behavior, fecundity and mortality is likewise determined according to equations (B7) through (B11), with the modification that in equations B8 and B10 the worker number is replaced by 1, assuming that egg-laying workers also have to provide for their own brood.

### Colony turnover and social structure

We work with a population of 1000 colonies for each simulation. Each colony is initialized with a single mated queen and no workers. A colony goes extinct once all its members have died. In this case, a new colony-founding queen is recruited at random from the brood of surviving colonies. The probability that a focal colony *k* with brood size *B_k_* supplies the new queen is 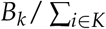, where *K* is the set of live colonies. A mate for the new queen is drawn similarly.

We investigate social structure by applying three “social scenarios” regarding the fate of a colony once its founding queen died: (i) monogyny, (ii) serial polygyny, and (iii) colony inheritance. In the monogyny scenario, the remaining brood develops into sexual offspring while the remaining workers, if they are not sterile, lay haploid eggs. Once the workers and sexuals have died, the colony is extinct and will be replaced. In the serial polygyny scenario, the colony recruits a new queen from its own brood, provided there is brood, resulting in potentially immortal colonies that are queenright. Important to note is that with this setup the queen’s exposure to extrinsic mortality is the same as under the monogyny scenario and no conflict over resources between queens emerges. We stress this because it has been argued that queens in polygynous species have relatively short lifespans because they are exposed to higher levels of extrinsic mortality than queens from monogynous species (Keller and Genoud 1997). Finally, in the colony inheritance scenario, a random worker takes over as new queen, and from then on she will express the “queen-part” of her genome. This scenario resembles primitively eusocial wasps such as paper wasps, where worker “queuing” for the chance to reproduce is believed to have been driving the evolution of long lifespans during the evolution of eusociality (Alexander et al. 1991; Bourke 2007; Hughes and Strassmann 1988).

### Simulation settings and software

Each simulation runs for 20, 000 discrete time steps, sufficient for (quasi-) evolutionary equilibrium to be reached. For each parameter setting we ran 50 replicate simulations. The code has been written in C++ (Stroustrup 2000) and compiled with GCC 5.1.0. To analyze and visualize the results we used R-statistical software (R Core Team 2017) and the packages pash (Schöley et al. 2017), ggplot (Wickham et al. 2016), phytools (Revell 2012), gridExtra (Auguie and Antonov 2017) and dplyr (Wickham et al. 2017).

