## Supplemental material for "Eusociality and the evolution of aging in superorganisms"

**Supplementary Material**

<u>*Simulation model parameter space*</u>

After an initial exploration of the parameter space, some parameters were held constant for all simulations presented in the manuscript: the number of colonies ($N$, Table 1) was kept constant at 1.000. Larger population sizes did not change the results but exponentially increased computation time. The simulated time steps used are 20.000 for all simulations, at 20 age classes this represents >1000 generations. Figure S1 shows that after ~10.000 time steps simulation reached an equilibrium where mutations and selection are balanced and changes to the evolved survival patterns are minimal. Longer simulation time do not change the results. For all simulations we used 20 age classes for both queens and workers ($\omega$ = 20, Table 1) resulting in 40 age specific genes that code for caste and age specific survival ($\varphi$ = 40). Simulations using 50 age classes ($\omega$ = 50) did not show qualitative differences, while with lower age class counts differences between queens and worker become smaller, because initial survival is important. Initial survival probabilities used at the start of each simulation for queens and workers were high and similar across all age classes ($a_0$ = 0.8, leading to a constant age specific survival probability across castes of 0.96 after the logistic transformation (eq. B1). Decreasing the initial survival probabilities leads to an increasing number of extinct populations where all colonies in the simulation die and no new queens are produced to fill extinct colonies. The mutation rate was kept constant for all simulations presented here ($m$ = 0.001). Simulations with lower mutation rates use more time steps to reach equilibrium, while larger rates lead to increased numbers of extinct populations if early mutations lead to decreased survival in the early age classes and selection cannot remove detrimental mutations. The parameter productivity ($r$, Table 1) controls the amount of resources foraged by one individual in one time step, determining the foraging success. Values for productivity used in the simulations lie between one and three, as otherwise colonies become very large, exponentially increasing the computational time needed for running the simulations. In the monogynous scenario low values of $r$ led to a strong decline of evolved queen lifespan with increasing extrinsic mortality because a reduced offspring production and small colony sizes force queens to forage (leading to exposure to extrinsic mortality), as a consequence deleterious mutation can accumulate in older age classes (Fig. S2, S3). Two parameters control the fecundity of queens and workers independently (parameters: $f_q$, $f_w$, Table 1). We tested different levels of fecundity for both queens and workers typically varying values between zero and six. As with the productivity parameter, increasing the fecundity leads to the production of more individuals and consequently larger colony sizes.

*Parameters that control for mutational effects*

Parameters that control the effects of mutations were kept constant (alpha, $\alpha = 0$, beta, $\beta = 0$, eta, $\eta = -0.2$, sigma, $\sigma = 0.08$, Table 1) for all simulations presented in the main text but we also tested alternative mutational effects that assume correlations between queen and worker phenotypes, or between age-classes. Transcriptome studies show similar genetic pathways are activated in queen and worker castes (Harrison et al. 2015) pointing to less independence of queen and worker phenotypes as so far assumed. We decreased the independence of worker and queen phenotypes by extending the mutational effect of a mutation that affects an age class in the worker phenotype to also affect the same age class of the queen phenotype with reduced effect size (parameter: beta, $\beta$), therefore increasing the correlation between queen and worker phenotypes. Less independent phenotypes generally lead to a reduction in phenotypic divergence (Fig. S4). In the monogynous scenario queens are slightly shorter lived while worker live longer as compared to settings with independent ageing phenotypes. When the correlation was high queen and workers showed similar lifespans when not being exposed to extrinsic mortality. Additional extrinsic mortality then drives the divergence of queen and worker phenotypes as worker lifespans decrease sharply and queen lifespan stays constant or decreases slightly. Also here sterile workers are short lived across different levels of extrinsic hazard while queen lifespans were slightly reduced when the correlations between phenotypes was strong. In the polygynous and the colony inheritance scenarios correlations between queen and worker phenotypes have a reduced effect since queen and worker lifespan divergence is generally less pronounced in these scenarios.

Alternatively age specific genes within a caste may be less independent than assumed so far. By increasing the effect of an age specific mutation to neighboring age classes with diminishing effect size (parameter: alpha, α) we decreased the independence of age specific genes within a caste. Across all scenarios increasing correlations between age classes lead to the evolution of longer lifespans of both queen and worker phenotypes (Fig. S5). Especially in the monogynous scenario queen (and worker in simulations without any extrinsic mortality) survival is maximal through all age classes and shows no decline with increasing extrinsic mortality, while worker lifespan decreases sharply along extrinsic mortality, maximizing the lifespan ratio. The lifespans of sterile workers were shorter than queen lifespans but due to correlations between age specific genes longer as if assuming independence of age specific genes. For the other scenarios queen and worker lifespans were increased but the observed response to increasing extrinsic mortality did not change when compare to results obtained from independent age specific genes. For the polygynous and the colony inheritance scenarios we found that the variance in evolved lifespans for both queens and workers between simulations with the same parameter setting increased (Fig. S5).

*Simulation model assumptions*

In the simulation model, we kept the genetics of aging as simple as possible by implementing a version of the mutation accumulation theory of aging (Medawar 1952). This model does not assume trade-offs between different age-specific physiological functions and is inherently more “neutral” than the antagonistic pleiotropy (Williams 1957) or disposable soma (Kirkwood 1977) theories of aging that postulate additional selection pressures on age-specific allocation of resources to reproduction vs. survival. We felt it is important to first explore the limits of a model with a minimal set of assumptions and selection pressures; otherwise, it might be too difficult to judge the relative importance of multiple simultaneously operating selection pressures. The implementation of trade-offs between physiological function could broaden view on the evolution of lifespan in eusocial systems and potentially explain the extreme lifespan ratios found in some species. However, this will require careful thinking, because the nature of trade-offs is especially complex in social insects because they live between two “levels of individuality”. Although selection experiments on solitary organisms have shown that lifespan extensions typically lead to a reduction in fecundity (Stearns et al. 2000) (but see (Maklakov et al. 2017)), trade-offs between fecundity and survival in social insects seem to disappear or may change in response to the social environment (Kramer et al. 2015; Negroni et al. 2016; Oettler and Schrempf 2016).

Monogyny and polygyny evoke different selection pressures on queen lifespans, resulting in shorter average queen lifespans in polygynous species(Keller and Genoud 1997). To keep processes in the models as similar as possible we chose to implement serial polygyny instead of allowing for multiple queens in one colony at the same time. Implementing multiple queens per colony requires further modelling assumptions such as the allocation of resources between different queens as well as allocation decisions on the colony level between investing in queens or workers. Serial polygyny is therefore the simplest way to account for the fact that queen death does not lead to the death of the entire colony in polygynous societies and besides avoiding potential allocation conflicts between queens results should not change if two queens exist in parallel or in succession.

*Supplemental information Figure 1*

We used phylogenetic ANOVAs (Garland et al. 1993; Revell 2012) to correct for the phylogeny and to compare lifespans between monogynous and polygynous species. We constructed a phylogeny from published phylogenies of social Hymenoptera (Schmitz and Moritz 1998; Danforth 1999; Costa et al. 2003; Arévalo et al. 2004; Moreau et al. 2006; Schultz and Brady 2008; Ward et al. 2016; Sparks et al. 2019) that contains 47 species of the 49 species in Figure 1. Because no information on branch lengths could be assigned to our composite phylogeny we set all branch lengths to the same value (0.1) and transformed the tree to an ultrametric tree before analysis. Table S1 shows the species used in the analysis.

### Supplementary Tables & Figures

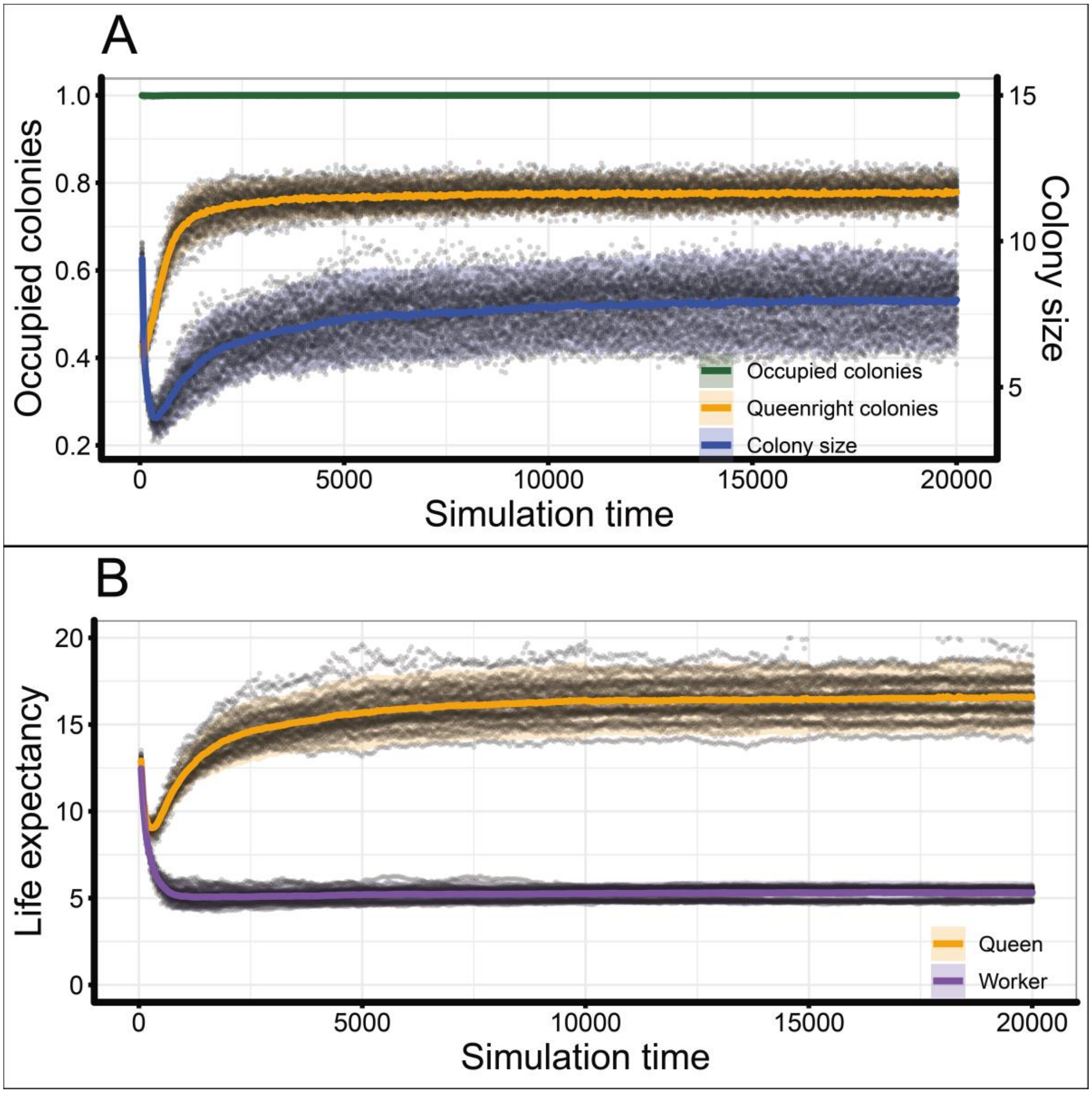


**Fig. S1.** Overview of 50 simulation replicates for the monogyny scenario with sterile workers. Panel A shows the proportion of occupied colonies (1000 colonies per population) in the simulations, as well as the proportion of momogynous colonies where the queen is alive. The blue curve indicates the mean number of workers per colony per simulation. Panel B shows the evolution of life expectancy at birth for workers and queens. For panels A and B gray dots indicate individual simulation runs while the lines and bands represent the mean and two standard deviations across all simulations. Parameter settings: $h_e = 0$, $r = 1.0$, $f_q = 5.0$, $f_w = 0$, $\alpha = \beta = 0$.

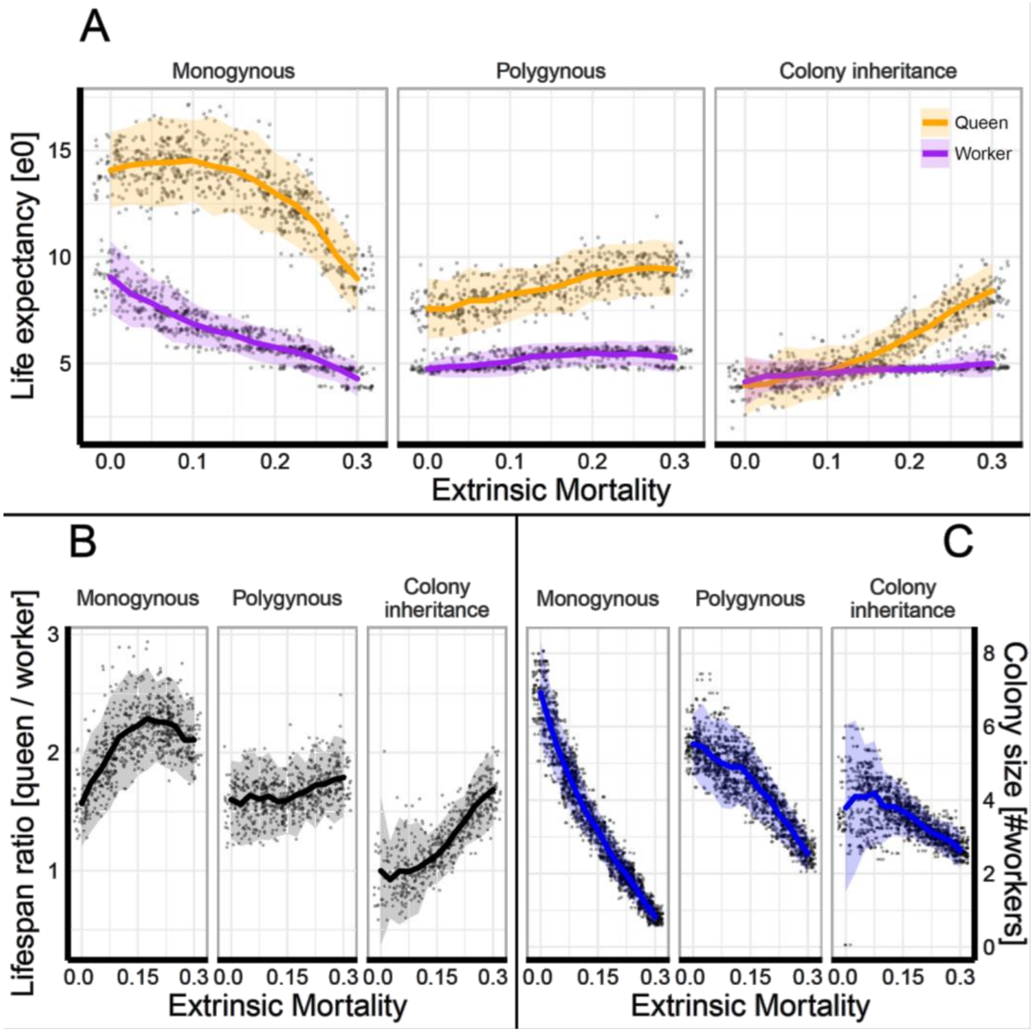


**Fig. S2.** The effect of extrinsic mortality and social structure on the evolution of queen and worker lifespan. Results of 50 simulations each for thirteen different levels of extrinsic mortality ($\boldsymbol{h_e}$) and three different scenarios (1950 simulations). Gray dots indicate the mean values from individual simulation runs, the solid lines represent means across 50 simulations and the shaded areas represent two standard derivations. Panel A shows evolved queen and worker life expectancies (e0) for the different scenarios (social structure) and levels of extrinsic mortality. Panel B shows the resulting lifespan ratio between queen and worker life expectancy. Panel C shows the evolved average colony sizes (number of workers) in the simulations. Parameters: productivity, $r = 1$, queen fecundity, $\boldsymbol{f_q} = 3$, worker fecundity, $\boldsymbol{f_w} = \mathbf{3}$.

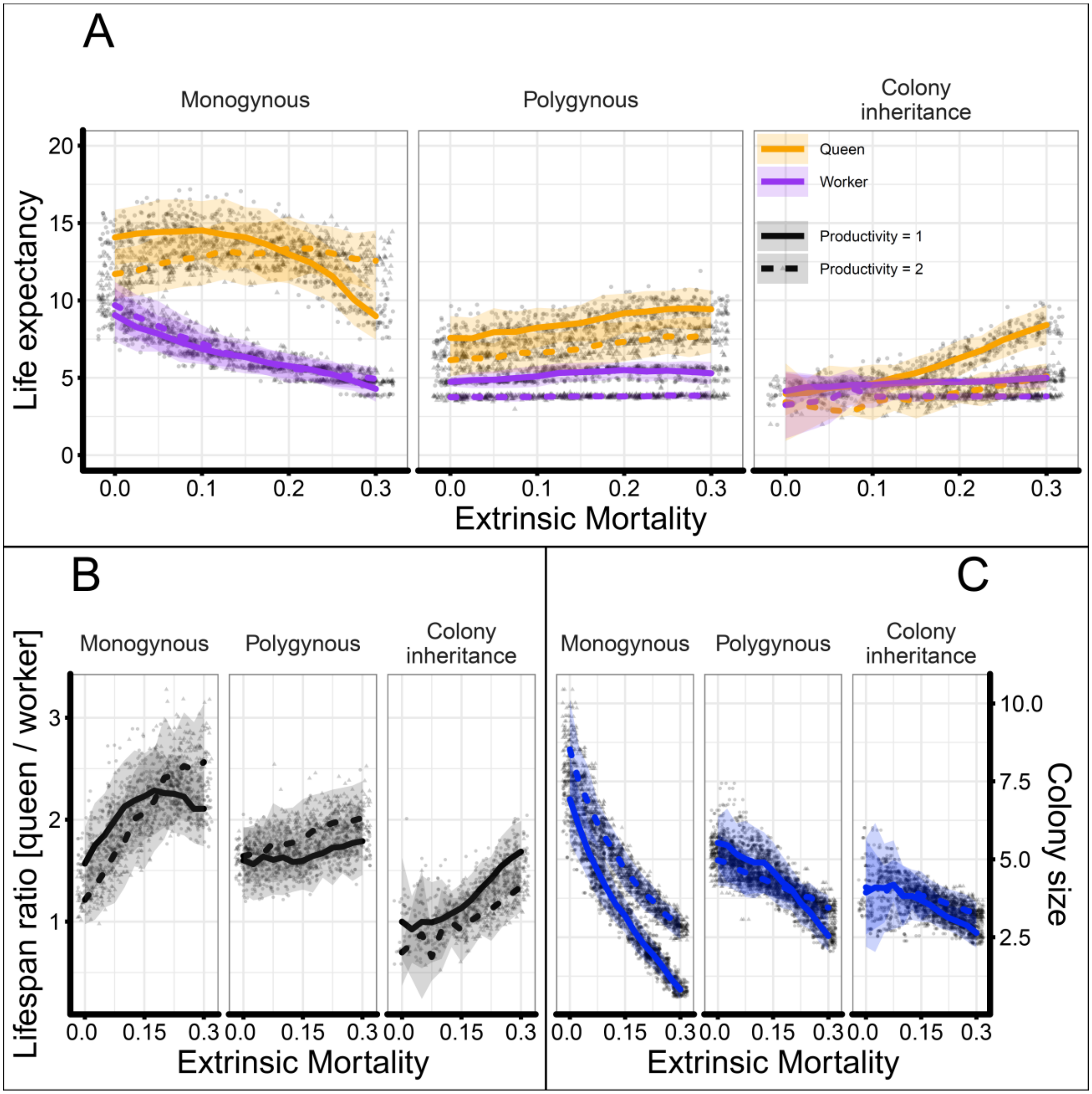


**Fig. S3.** The effect of productivity on the evolution of queen and worker lifespans. Results of 50 simulations each for thirteen different levels of extrinsic mortality ($\boldsymbol{h_e}$) and three different scenarios. Gray dots indicate the mean values from individual simulation runs, the solid (dashed) lines represent means across 50 simulations with low productivity (high productivity) (parameter: $r$ ), which represents the foraging success. The shaded areas represent two standard derivations. Panel A shows evolved queen and worker life expectancies at birth for the different scenarios (social organisation) and levels of extrinsic mortality. Panel B shows the resulting lifespan ratio between queen and worker life expectancy. Panel C shows the evolved average colony sizes (number of workers) in the simulations. Parameters: productivity, $r$ = {1, 2}, queen fecundity, $\boldsymbol{f_q}$ = 3, worker fecundity, $\boldsymbol{f_w}$ = **3**, $\alpha$ = 0, $\beta$ =0.

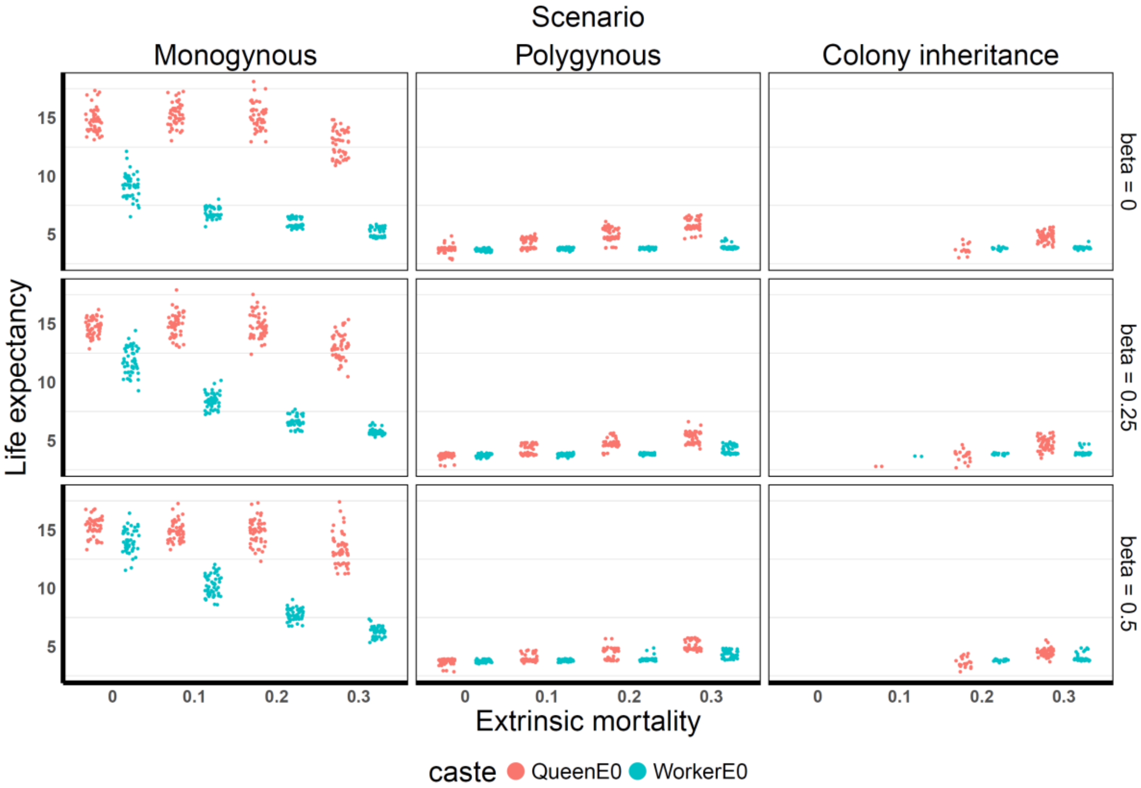


**Fig. S4**

Effect of phenotypic independence (parameter: beta, β). Parameter settings: productivity, $r$ =1, queen fecundity, $f_q$ = 6, worker fecundity, $f_w$ = 3, alpha, α= 0, beta, β={0,0.25,0.5}, sigma, σ=0.08. Red points represent queens, blue workers. Increasing values of beta increase correlations between queen and worker castes as mutational effects between castes are correlated. As a result the independence between castes is reduced and the divergence in lifespan between queens and workers is reduced

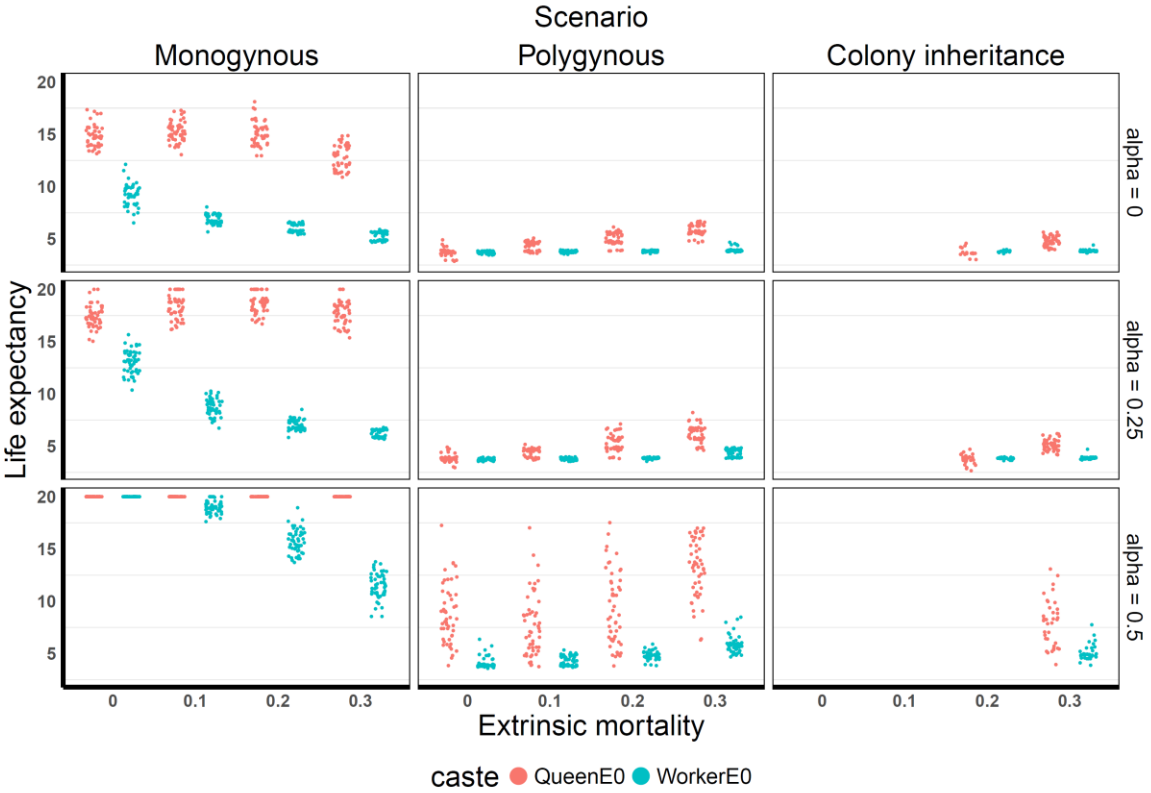


**Fig. S5**

Effect of age class independence (parameter: alpha, α). Parameter settings: productivity, $r$ =1, queen fecundity, $f_q$ = 6, worker fecundity, $f_w$ = 3, $\alpha$ = {0,0.25,0.5}, $\beta$ = 0, $\sigma$ = 0.08. The points represent mean intrinsic life expectancy per simulation, red points present queens, blue workers. Increasing values of $\alpha$ (0-1) increase correlations between age classes, as mutations affect genes coding for neighboring age classes within a caste or phenotype, rendering age classes less independent.

**Table S1** Species used in figure 1 (data from Kramer and Schaible 2013).

| Family | Genus | Species | Social structure |
|---|---|---|---|
| Apidae | Apis | mellifera | Monogynous |
| Apidae | Bombus | melanopygus | Monogynous |
| Apidae | Bombus | terrestris | Monogynous |
| Apidae | Melipona | favosa | Monogynous |
| Apidae | Melipona | quadrifasciata | Monogynous |
| Apidae | Tetragonisca | angustula | Monogynous |
| Formicidae | Acromyrmex | octospinosus | Monogynous |
| Formicidae | Aphaenogaster | rudis | Monogynous |
| Formicidae | Atta | cephalotes | Monogynous |
| Formicidae | Atta | colombica | Monogynous |
| Formicidae | Atta | sexdens | Monogynous |
| Formicidae | Diacamma | rugosum | Monogynous |
| Formicidae | Formica | fusca | Monogynous |
| Formicidae | Harpagoxenus | saltator | Monogynous |
| Formicidae | Lasius | flavus | Monogynous |
| Formicidae | Lasius | niger | Monogynous |
| Formicidae | Messor | semirufus | Monogynous |
| Formicidae | Myrmecia | gulosa | Monogynous |
| Formicidae | Myrmecia | nigriceps | Monogynous |
| Formicidae | Myrmecia | vindex | Monogynous |
| Formicidae | Paraponera | clavata | Monogynous |
| Formicidae | Pogonomyrmex | badius | Monogynous |
| Formicidae | Pogonomyrmex | salinus | Monogynous |
| Formicidae | Rhytidoponera | purpurea | Monogynous |
| Formicidae | Temnothorax | lichtensteini | Monogynous |
| Formicidae | Temnothorax | nylanderi | Monogynous |
| Formicidae | Trachymyrmex | septentrionalis | Monogynous |
| Halictidae | Halictus | marginatus | Monogynous |
| Halictidae | Lasioglossum | marginatum | Monogynous |
| Vespidae | Polistes | chinensis | Monogynous |
| Vespidae | Polistes | cinerascens | Monogynous |
| Vespidae | Polistes | fuscatus | Monogynous |
| Vespidae | Polistes | versicolor | Monogynous |
| Vespidae | Ropalidia | fasciata | Monogynous |
| Vespidae | Vespa | orientalis | Monogynous |
| Vespidae | Vespa | simillima | Monogynous |
| Vespidae | Vespa | tropica | Monogynous |
| Formicidae | Monomorium | cyaneum | Polygynous |
| Formicidae | Monomorium | ergatogyna | Polygynous |
| Formicidae | Monomorium | pharaonis | Polygynous |
| Formicidae | Monomorium | trageri | Polygynous |
| Formicidae | Monomorium | viride | Polygynous |
| Formicidae | Myrmica | rubra | Polygynous |
| Formicidae | Platythyrea | punctata | Polygynous |
| Halictidae | Halictus | ligatus | Polygynous |
| Vespidae | Mischocyttarus | atramentarius | Polygynous |
| Vespidae | Mischocyttarus | cerberus | Polygynous |
| Vespidae | Mischocyttarus | extinctus | Polygynous |
| Vespidae | Ropalidia | marginata | Polygynous |
